# Nucleic acid delivery of immune-focused SARS-CoV-2 nanoparticles drive rapid and potent immunogenicity capable of single-dose protection

**DOI:** 10.1101/2021.04.28.441474

**Authors:** Kylie M. Konrath, Kevin Liaw, Yuanhan Wu, Xizhou Zhu, Susanne N. Walker, Ziyang Xu, Katherine Schultheis, Neethu Chokkalingam, Jianqiu Du, Nicholas J. Tursi, Alan Moore, Mansi Purwar, Emma L. Reuschel, Drew Frase, Matthew Sullivan, Igor Maricic, Viviane M. Andrade, Christel Iffland, Kate E. Broderick, Laurent M. P. F. Humeau, Trevor R.F. Smith, Jesper Pallesen, David B. Weiner, Daniel W. Kulp

## Abstract

Antibodies from SARS-CoV-2 vaccines may target epitopes which reduce durability or increase the potential for escape from vaccine-induced immunity. Using a novel synthetic vaccinology pipeline, we developed rationally immune focused SARS-CoV-2 Spike-based vaccines. N-linked glycans can be employed to alter antibody responses to infection and vaccines. Utilizing computational modeling and comprehensive in vitro screening, we incorporated glycans into the Spike Receptor-Binding Domain (RBD) and assessed antigenic profiles. We developed glycan coated RBD immunogens and engineered seven multivalent configurations. Advanced DNA delivery of engineered nanoparticle vaccines rapidly elicited potent neutralizing antibodies in guinea pigs, hamsters and multiple mouse models, including human ACE2 and human B cell repertoire transgenics. RBD nanoparticles encoding wild-type and the P.1 SARS-CoV-2 variant induced high levels of cross-neutralizing antibodies. Single, low dose immunization protected against a lethal SARS-CoV-2 challenge. Single-dose coronavirus vaccines via DNA-launched nanoparticles provide a platform for rapid clinical translation of novel, potent coronavirus vaccines.

## Introduction

Severe Acute Respiratory Syndrome 2 (S3RS-CoV-2) virus is responsible for Coronavirus disease 2019 (COVID-19) in over 144 million people and 3.0 million deaths as of April 22^th^ 2021[1, 2]. The Spike(S) glycoprotein studs the surface of Coronaviruses virions and its receptor-binding domain (RBD) binds host cell receptors to mediate viral entry and infection[3, 4]. Greater than 90% of COVID-19 patients produce neutralizing antibodies (nAbs)[5] and RBD-directed antibodies often comprise 90% of the total neutralizing response[6]. RBD-directed antibodies can correlate with neutralizing activity[7–9] and ~2,500 antibodies targeting the SARS-CoV-2 spike have been described to date[10, 11]. This highlights the importance of eliciting neutralizing antibodies targeting the RBD by vaccination.

Rational SARS-CoV-2 vaccine design should be informed by spike protein conformation dynamics, the sites of vulnerability and mutations that cause potential vaccine escape. The S trimer has >3,000 residues creating a vast array of epitopes and is targeted by both neutralizing and non-neutralizing antibodies(non-nAbs)[12–15]. Measures of RBD binding do not always correlate with neutralization due to presence of non-nAbs, which have the potential to cause antibody-dependent enhancement[16–18]. In the context of HIV, influenza and MERS-CoV, significant effort over the last few decades has focused on creating immunogens that minimize non-neutralizing epitopes[19–26]. Since the initial outbreak of SARS-CoV-2, significant headway has been made in identifying neutralizing epitopes, especially with regards to the RBD; however, study of immunodominant, non-neutralizing epitopes has lagged[14, 27, 28]. Vaccine immunogens should be developed with these key findings in mind.

Glycosylation is an important post-translational modification in viral pathogenesis serving versatile roles including host cell trafficking and viral protein folding[29]. Mutations introducing potential N-linked glycosylation sites (PNGS) [30] in other viruses such as HIV and influenza have contributed to immune escape[31–34]. Structure-based vaccine design efforts have been employed to add exogenous PNGS to block non-neutralizing sites and focus the immune response to neutralizing sites[20, 26, 35, 36]. These approaches have not been widely applied to SARS-CoV-2 vaccine development. Here, we develop an advanced structural algorithm for optimizing PNGS into the SARS-CoV-2 RBD to focus the immune response and enhance neutralizing responses targeting the Receptor Binding Site epitope(RBS).

Vaccine potency is important for an effective immunological response. Self-assembling, multivalent nanoparticle immunogens (or nanovaccines) enhance the B cell activation and concomitant humoral responses, kinetics of trafficking to the draining lymph nodes and uptake by dendritic cells and macrophages[37–40]. SARS-CoV-2 nanovaccines developed as recombinant proteins can be difficult to clinically translate due to arduous purification and manufacturing processes, and further do not tend to activate CD8+ T cells[38]. In contrast, vaccine antigens encoded into a DNA plasmid can be delivered directly *in vivo*. We recently demonstrated the speed of DNA vaccine translation by developing a DNA-encoded full-length spike immunogen for clinical evaluation in 10 weeks[41]. DNA is easily mass produced, temperature stable, not associated with anti-vector immunity and can be rapidly reformulated for circulating variants, making it a key pandemic vaccine technology. We recently developed a DNA-launched nanoparticle platform (DLNP) for *in vivo* assembly of nanoparticles which drive rapid and strong B cell immunity and uniquely produce strong CD8+ T cells[38]. Here, we present a SARS-CoV-2 DLNP which has enhanced immunity in multiple animal models and is capable of single shot protection against lethal challenge. The single shot, low dose regimen reduces the overall amount of necessary product, medical personnel, and time in the clinic, rendering the product more scalable to a global scope including resource limited settings. The high potency and rapid developability of the DLNP platform can also enable quick generation of booster vaccines for newly emergent variants. To this end, we developed a SARS-CoV-2 DLNP encoding P.1 mutations and demonstrate it is highly immunogenic.

## Results

### Mapping antigenic effects of N-linked glycans on the RBD

To assess the feasibility of adding N-linked glycans to alter antibody responses to RBD (Figure 1A), we built an advanced structural algorithm called Cloaking With Glycans (CWG) for modeling every possible glycan on the RBD (Figure 1B). The PNGS positions were filtered if the asparagine had low solvent accessibility or high clash score (Figure 1C). Next, we surveyed energetics of naturally occurring glycans (Figure S1A, S1B) and employed glycan energy filters for our designed glycan positions, as well as filters for protein folding energies and structural considerations (see methods). This process led to the identification of 43 out of 196 positions for experimental characterization (Figure 1C,1D).

**Figure 1:**
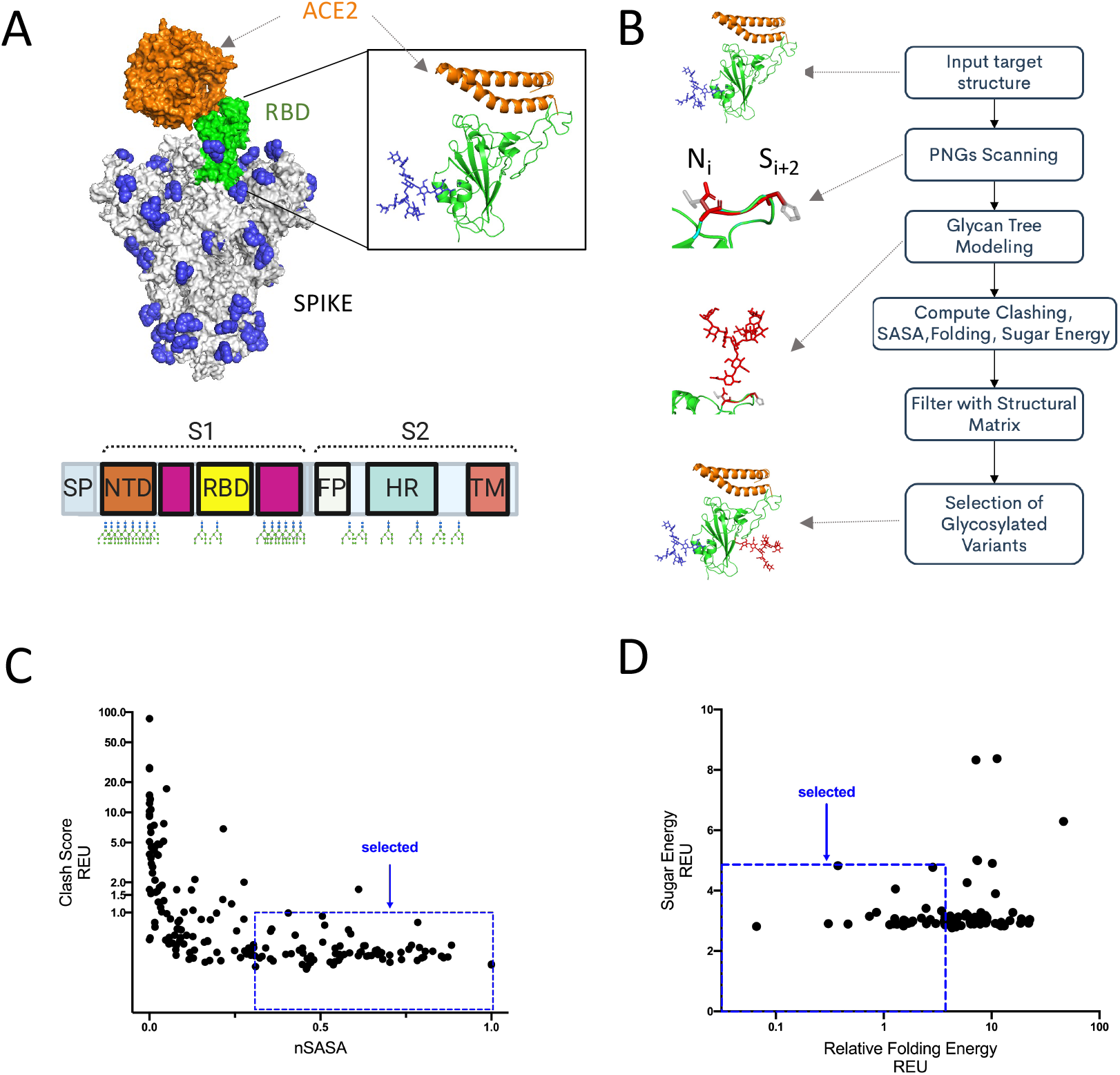
CWG algorithm to identify sites amenable to glycosylation. (A) SARS-CoV-2 spike trimer (grey) decorated with native glycans (blue) with one RBD in the up state (green) binding to ACE2 (orange) and detailed cartoon representation of RBD with native glycan bound to ACE2. Schematic of wild type glycan distribution across the entire spike. (B) CWG pipeline for assessing PNGS on the RBD. (C) Rosetta scores of glycosylated RBDs for normalized solvent accessible surface (SASA) and residue clash score (fa_rep of sugar residues). (D) Protein folding (total Rosetta score) vs Glycan score (fa_rep of sugar and protein) for each of the glycosylated RBDs selected in (C). Selection criteria shown as dashed lines in (C) and (D).

To better understand single glycan mutants (Figure 2A), we produced each variant and measured biophysical and antigenic profiles. We synthesized and screened the glycan variants for expression and binding to ACE2 in a high-throughput, small-scale transfection format and downselected to 22 variants for further evaluation (Figure 2B). To characterize the antigenic properties of the glycan variants, we utilized 14 RBD-directed nAbs, 2 Abs with inconsistent neutralization[42–45], and 6 non-nAbs[10, 12, 13, 15, 44–51]. Most nAbs target epitopes in the RBS (RBD-A, RBD-B, RBD-C)[10] and some target outside the RBS[27] (RBD-D, RBD-E and RBD-F) (Figure 2C). In general, we sought to identify glycans that do not interfere with nAb binding and block non-nAbs. The reactivity of our set of antibodies to each glycan mutant was determined by SPR and ELISA (Figure 2D,2E, Table S1). We observed reduced binding of neutralizing RBD-A, RBD-B, RBD-C or RBD-D antibodies in the presence of glycans at residues 441, 448, 450, 458 and 481, suggesting these could be potential vaccine escape mutations because they still bind human ACE2. In addition, glycans at positions 337, 344, 354, 357, 360, 369, 383, 448, 450, 516 and 521 show dramatically reduced binding to non-nAb(s). We did not observe effects on binding to our antibody panel for glycans at 518, 519 and 520. We noticed similar antigenic patterns in glycan positions that reduce binding to some of the non-nAbs as well as nAbs in RBS-E and RBS-F, suggesting there is overlap in these nAb and non-nAb epitopes. In sum, our experimental screening exhaustively evaluated the effect of N-linked glycans on the expression and antigenic profile of the RBD.

**Figure 2:**
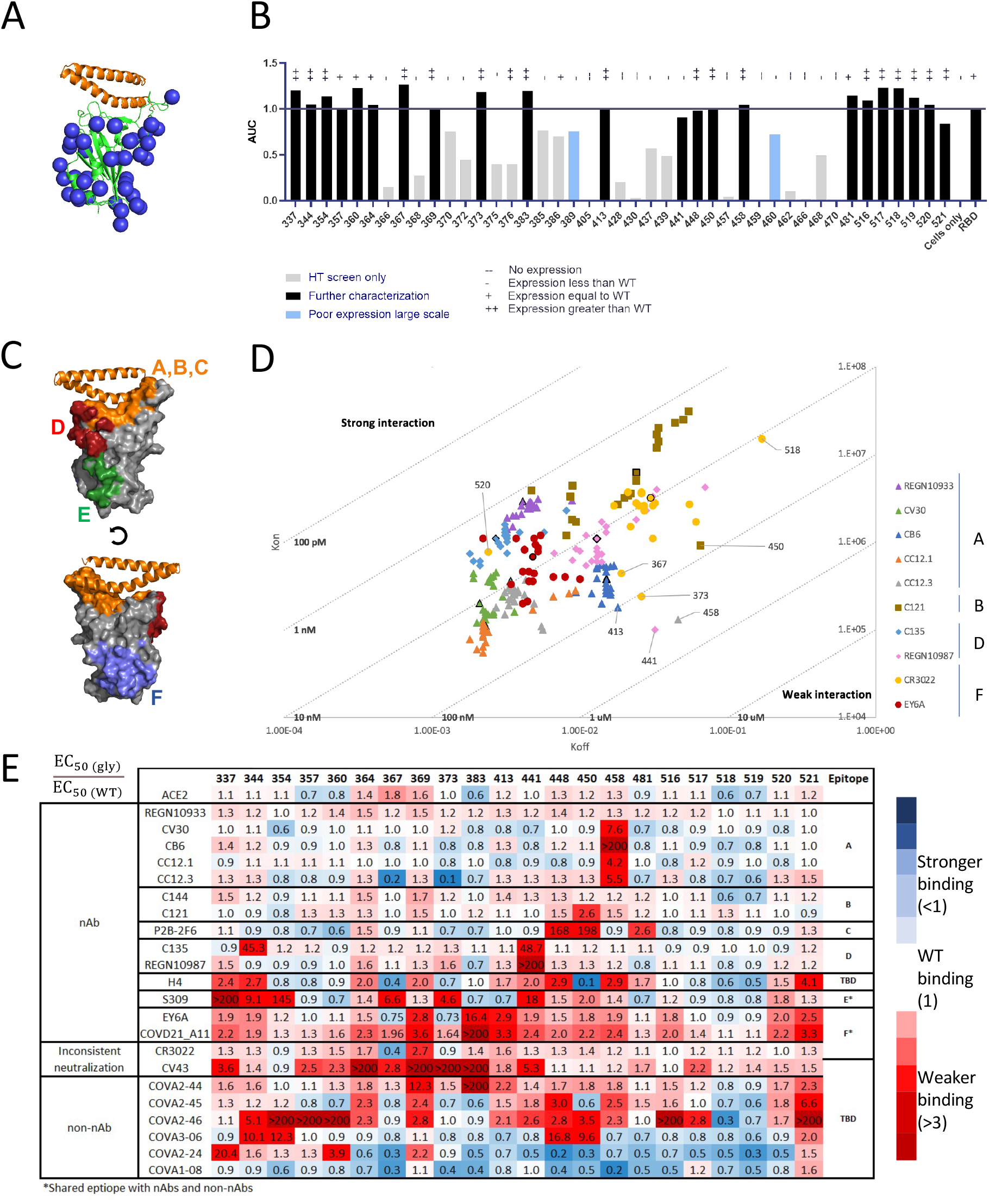
In vitro characterization of single glycan variants of RBD. (A) Model of selected glycan sites (blue spheres) on the RBD (green cartoon) interacting with ACE2 binding helices (orange cartoon). (B) Small scale screen of selected variants binding to ACE2 in Area Under the Curve from ELISA binding curves and normalized to WT binding (bars), qualitative expression from Western Blot represented as +/- symbols above the bars. (C) Neutralizing epitopes mapped on RBD structure with RBD in grey surface and ACE2 binding helices in orange, surface patches are color according to: RBD-A,B,C are in orange; RBD-D in red; RBD-E in green, RBD-F in blue. (D) SPR binding kinetics of single glycan variants to a panel of SARS-CoV-2 antibodies. (E) Relative binding as measured by ELISA EC50 ratio of glycan variants binding to WT RBD binding in a panel of neutralizing and non-neutralizing antibodies. Blue to Red coloring was done based on stronger or weaker binding relative to WT RBD.

### N-linked glycan decoration improves RBD directed immunity

We utilized our single glycan data to add sets of glycans to the RBD that maximally cover multiple non-neutralizing epitopes and preserve accessibility to RBS targeted neutralizing epitopes. To this end, we constructed a glycan distance map allowing design of three, five and eight glycan combinations which were experimental tested to determine if the sets could provide optimized antigenic profiles (Figure 3A, S2A, S2B, S2C). Two of the three glycan variants (g3.1 and g3.3) had heavily reduced binding to all antibodies in our panel. Eight glycan variants (g8.1, g8.2 and g8.3) had slightly reduced EC50 to nearly all the RBD neutralizing antibodies (Figure 3B). However, both g3.2 and g5.1 (harboring three and five glycans, respectively) bound well to nAbs and had reduced affinity for non-nAbs (Figure 3B). Since new non-nAbs may be identified in the future, we focused the remaining experiments on the more glycosylated variants (i.e. g5.1 over g3.2), since they are more likely to reduce accessibility to epitopes recognized by non-nAbs. We observed similar immunogenicity from BALB/c mice immunized with wild-type (WT) RBD or g5.1 RBD (Figure 3C,3D). To investigate the difference in specificity of the RBD elicited responses, we employed an ACE2 blocking assay[52] (Figure 3E). We observed that RBD g5.1 elicited significantly more ACE2-blocking antibodies than WT RBD, suggesting g5.1 is immune focusing antibodies to the RBS (Figure 3F, 3G). This data demonstrates that combinations of strategically selected glycans reduce the affinity of non-nAbs and focus immune responses to the neutralization-rich RBS or other epitopes of interest.

**Figure 3:**
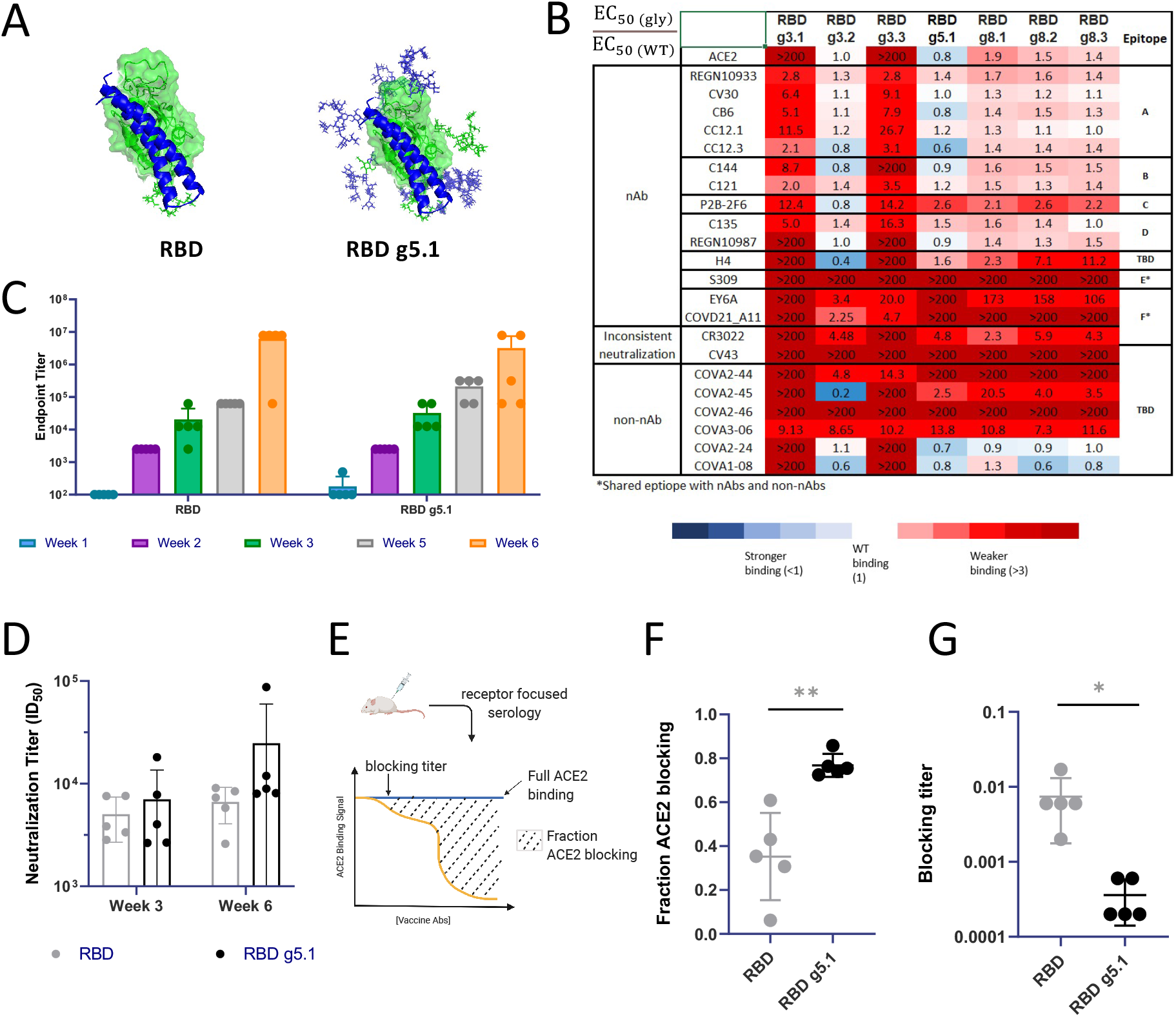
In vitro and in vivo antigenic profile of multiglycan RBDs. (A) Surface representation of RBD (green) bound to ACE2 (blue cartoons) with glycans (green for native and blue for designed in lines) for the WT RBD and RBD g5.1 constructs. (B) Relative binding as measured by ELISA EC50 ratio of glycan variants binding to a panel of neutralizing and non-neutralizing antibodies to WT RBD binding. Blue to Red coloring was done based on stronger or weaker binding relative to WT RBD. (C) Antibody binding titers and (D) pseudovirus ID50 neutralization titers from BALB/c mice immunized with 25μg of plasmids encoding WT RBD or RBD g5.1 at week 0 and 2. (E) ACE2 competition assay layout for measuring blocking of ACE2 interacting with RBD with RBS-directed antibodies from the sera of vaccinated mice. (F) Fraction of ACE2 binding blocked by antibodies in ACE2 competition assay and (G) Blocking titer measured as the first dilution of sera at which a reduction in ACE2 binding is observed. (unpaired two tailed Student t-test (F) p = 0.0020, (G) p = 0.0236).

### DNA-Launched nanovaccines amplify and accelerate immune responses

To develop multivalent vaccines, we genetically fused RBDs to a set of four different self-assembling scaffold proteins[38–40, 53] with a potent CD4-helper epitope(LS-3) to help enhance germinal center responses[54]. Tandem repeats of RBD have been shown to improve neutralization titers by 10-100 fold [55], thus we displayed dimers of RBDs on some of our self-assembling scaffolds as well. We engineered nanoparticles using our computational design methods[38], resulting in display of 7, 14, 24, 48, 60, 120 or 180 RBDs (Figure 4A). We rapidly screened 19 nanoparticles directly *in vivo* using a single mouse per construct at a single low dose of 2μg. We observed that 14 of the 19 nanoparticles were immunogenic (Figure 4B). Strikingly, rapid antibody responses were detected just 1 week after immunization with RBD g5.1 24mer, RBD 48mer and RBD g5.1 120mer (Figure 4B). In parallel, we expressed and purified nanoparticles *in vitro*. In contrast to wild type RBD multimers, which we could not purify and were not more immunogenic than RBD monomer (Figure S3A,S3B), we were able to purify nine glycan modified RBD multimers as assessed by size exclusion chromatography with multiangle light scattering (Figure 4B,4C). To further confirm assembly of the RBD g5.1 24mer, we employed structural analysis by cryo electron microscopy (cryo-EM) for RBD g5.1 24mer (Figure 4D, 4E). We conducted immunizations with selected constructs in BALB/c mice (n=5 or n=10) using a single low dose of 2μg (Figure 4F). RBD g5.1 24mer and 120mer both generated strong binding and neutralizing responses (mean ID_50_ of 3677 and 791, respectively) (Figure S3C). In C57BL/6 mice, we observed similar immunogenicity at 1μg and 5μg doses for select nanoparticles including improvements in CD8+ T cells (Figure S4A–S4E). RBD g8.2 7mer and RBD g8.2 24mer elicit similarly strong humoral responses when administered as purified protein nanoparticles (Figure S5). Strikingly, we observed strong binding and neutralizing responses in BALB/c mice immunized with 5μg RBD g5.1 24mer against the emergent South Africa (B.1.351), UK (B.1.17) and the Brazilian variant (P.1), indicating cross-reactivity and strong potential relevance against emerging variants (Figure 4G, S6A). As proof-of-concept for expanding this platform to emerging variants, we engineered P.1 RBD g5.1 24mers. Upon of BALB/c mice immunization with 2μg, we observed high binding and cross-neutralization titers (Figure 4H, 4I, S6B).

**Figure 4:**
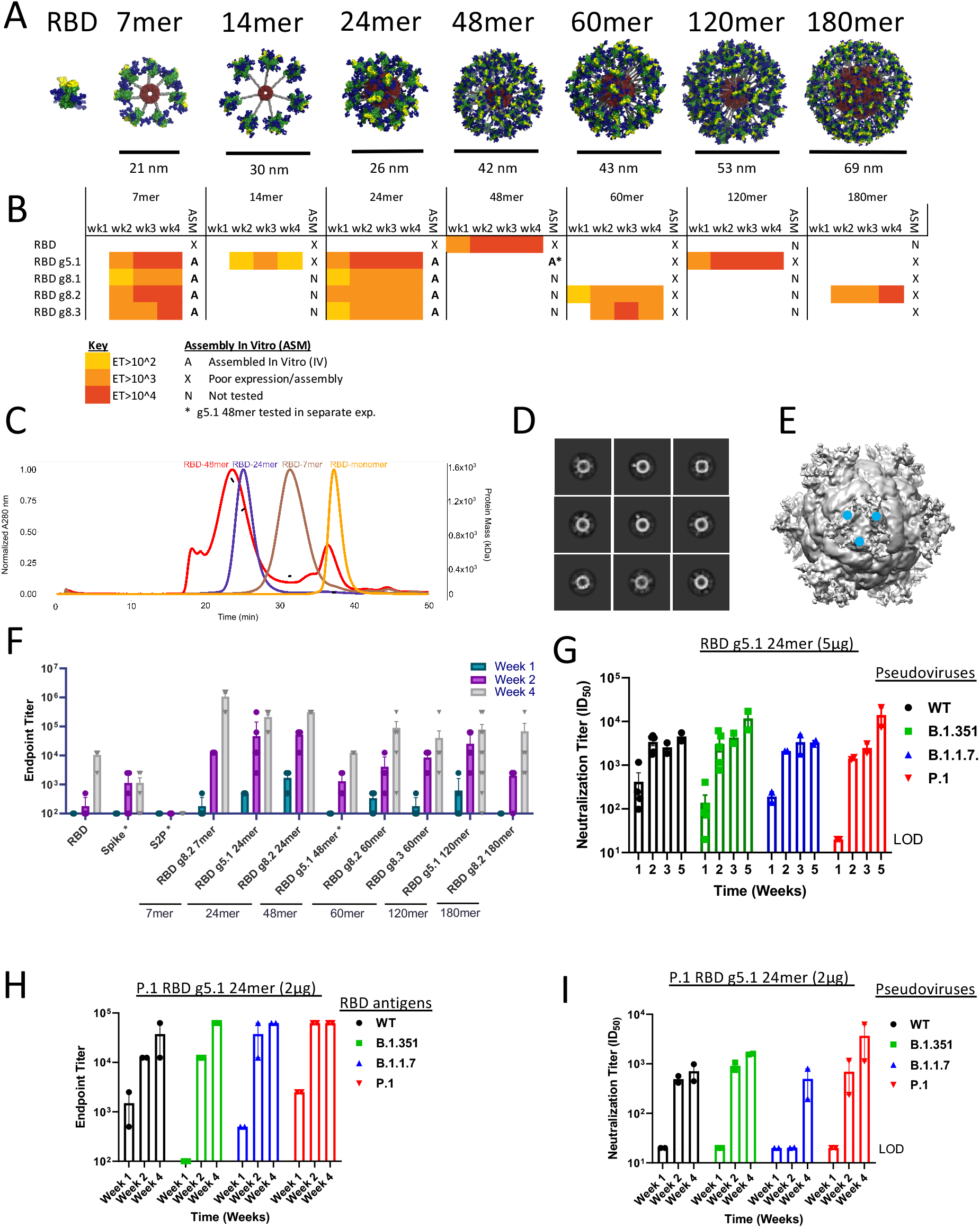
Immune focused RBD nanoparticle structure and immunogenicity. (A) Models of 8 different RBD nanovaccines. In each model, the coloring is as follows: RBS (yellow) on the RBD (green) coated with glycans (blue) fused with a glycine-serine linker (gray) to a nanoparticle scaffold (red). (B) Endpoint titers for a single BALB/c mouse immunized with once with 2 μg of plasmid encoding RBD nanoparticles by DNA-E.P. colored as indicated on the figure, in vitro expression and assembly of nanoparticles indicated in the ‘ASM’ column as either expressed/assembled (A), poor expression/assembly (X) or not tested (N). (C) Size-exclusion chromatogram and multiangle light scattering of RBD g5.1 multimers (Black line under each curve indicates molecular weight and correspond to the right y-axis). (D) 2D class averages showing RBDs decorating the RBD g5.1 24mer. (E) Cryo-EM density map of RBD g5.1 24mer at low threshold, the 24mer scaffold could be unambiguously determined (Figure S9), the flexible linker attachment points for the RBDs on the 24mer scaffold could be observed at low density threshold (blue dots) (F) Endpoint titers for expanded groups (n=5) of BALB/c mice immunized with 2μg of plasmid encoding RBD nanoparticles by DNA-E.P. (G) Pseudovirus neutralization of SARS-CoV-2 variants B.1(WT), B.1.351, B.1.7.1 and P.1 by sera from BALB/c mice immunized with 5μg RBD g5.1 24mer. (H) Endpoint binding titers developed in BALB/c mice immunized with 2μg of P.1 RBD g5.1 nanoparticle (I) Pseudovirus neutralization of SARS-CoV-2 variants by sera from BALB/c mice immunized with P.1 RBD g5.1

### Single dose of RBD nanoparticles affords protection in lethal challenge model

To examine the efficacy of the SARS-CoV-2 RBD nanoparticles with rapid seroconversion(RBD g5.1 24mer and RBD g5.1 120mer), we pursued a lethal challenge study (Figure 5A). B6.Cg-Tg(K18-ACE2)2Prlmn/J(K18-hACE2) mice express human ACE2 on epithelial cells including in the airway[56, 57] and can be infected with SARS-CoV-2 resulting in weight loss and lethality[58] providing a stringent model for testing vaccines[59]. Animals were vaccinated with a single shot of 5μg and 1μg of our nanovaccines in K18-ACE2 mice representing doses 5- and 25-fold lower than our standard DNA dose[41]. Prior to a blinded challenge, we examined immunogenicity at day 21 and observed pseudovirus neutralization titers prior to challenge in all vaccine groups (Figure S7). We also observed live SARS-CoV-2 virus neutralization titers above the limit of detection for all three nanoparticle groups with mean ID_50_ of 451, 1028, and 921 for RBD g5.1 120mer 1μg, 5μg and RBD g5.1 24mer 5μg, respectively, compared to a mean ID_50_ of 29 of RBD monomer (Figure 5B). The mice were infected with SARS-CoV-2 1×10^5^ PFU/mouse intranasally and monitored for signs of deteriorating health. We observed that mice immunized with nanovaccines had higher levels of protection from weight loss(Figure 5C). As expected, the naive group of animals reached 100% morbidity by day 6 and 1/10 animals survived in the RBD monomer group. In both RBD g5.1 120mer groups had 6/10 mice survived the challenge. Strikingly, immunization with RBD g5.1 24mer provided full protection from a lethal SARS-CoV-2 challenge (Figure 5D). All but one animal that survived had a live virus neutralization titer of >100 and 12/15 of the mice that succumbed to infection did not have appreciable neutralization titers (Figure 5E). We observed a significant correlation between live virus neutralization ID_50_ titer and body weight loss (Figure 5F). Viral replication was reduced in nasal turbinates, lung tissue and brain tissue for mice immunized with nanovaccines relative to RBD monomer or naïve animals (Figure 5G). Thus, the DNA-launched nanovaccines can generate potent immunity that provides protection from challenge with a single immunization at a low dose.

**Figure 5:**
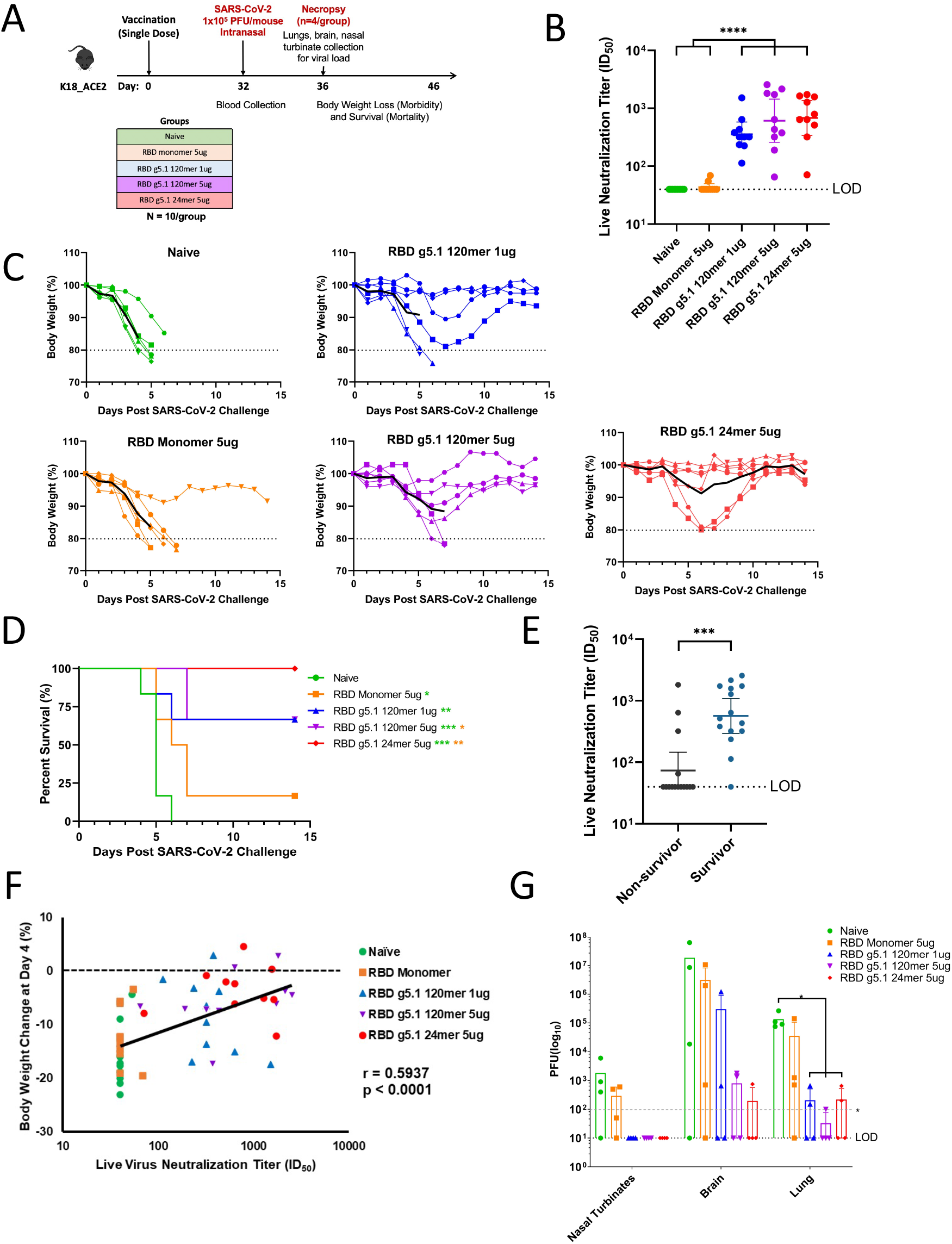
Lethal challenge of SARS-CoV-2 in rodent model. (A) K18 hACE2 lethal challenge study overview (B) SARS-CoV-2 live virus neutralization one day prior to challenge. **** p < 0.0001. (C) Weight loss of K18 hACE2 mice after SARS-CoV-2 challenge. (D) Kaplan-Meier curves representing survival of K18 hACE2 mice after SARS-CoV-2 challenge. (Mantel-Cox test vs. naïve: RBD monomer p = 0.0327, RBD g5.1 24mer p = 0.0006, RBD g5.1 120mer 1 μg p = 0.0087, RBD g5.1 120mer 5 μg p = 0.0006; vs. RBD monomer: RBD g5.1 120mer 5 μg p = 0.0426, RBD g5.1 24mer p = 0.0048) (E) Pseudovirus neutralization titers of surviving and non-surviving mice. (unpaired two-tailed Student t-test p = 0.0003) (F) Correlation between body weight change at day 4 post-challenge and pre-challenge live virus neutralizing titers (ID_50_). (G) Viral titers in nasal turbinates, brain and lung tissue at day 4 post challenge. (unpaired two-tailed Student t-test vs. naïve: RBD 120mer 1 μg p = 0.0110, RBD 120mer 5 μg p = 0.0109, RBD g5.1 24mer p = 0.0110). LOD for this assay (lower dashed line) is lower than the LOD reported elsewhere (top dashed line).

### Enhanced immune responses to nanovaccines in translational vaccine models

One major challenge for the clinical translation of vaccines is preclinical modeling of human antibody responses to immunogens. OmniMouse^®^ have humanized immunoglobulin loci-transgenic with human V, D and J gene segments[60]. As a proof-of-concept, we immunized OmniMouse^®^(n=3) with three different SARS-CoV-2 nanoparticle vaccines and measured increasing RBD-specific human antibodies in serum (Figure 6A,6B). Most mice produced high titers of IgG and a few had robust IgM titers (Figure S8A,S8B). We observed potent and specific neutralization in all three groups at weeks 6 and 8 (Figure 6C, S8C). Thus, the SARS-CoV-2 nanoparticle platform can be employed in transgenic mice and induce human SARS-CoV-2 neutralizing antibodies.

**Figure 6:**
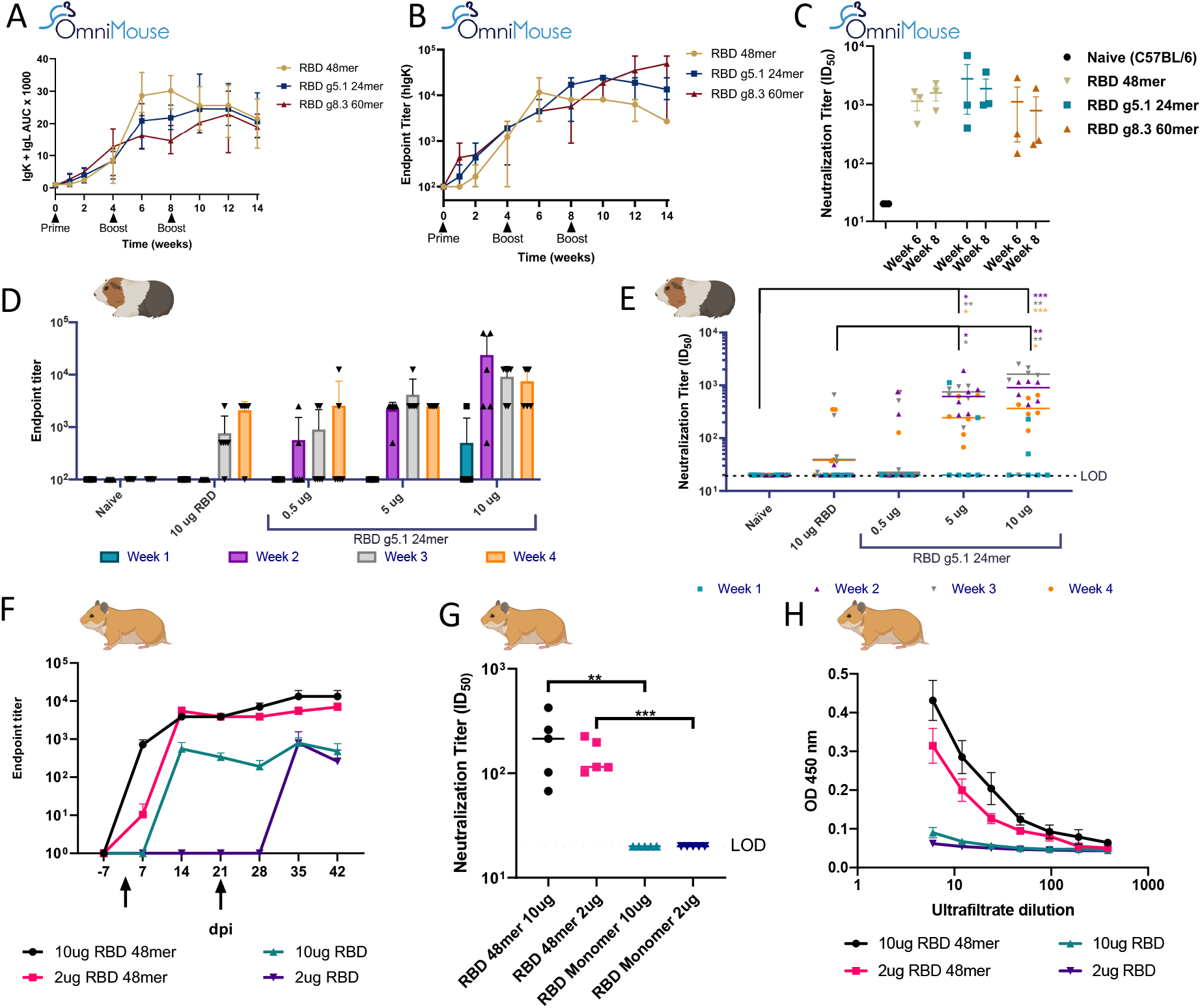
Humoral responses to nanovaccines in OmniMouse^®^, Guinea Pigs and Hamsters. (A) Human antibody titers from OmniMouse^®^ immunized three times four weeks apart with 25μg of DNA encoding RBD nanoparticles as measured by combined AUC from ELISA curves with human IgK and human IgL secondaries. (B) as measured by endpoint titer using human IgK and human IgL secondaries (C) Pseudo virus neutralization titers at week 6 and week 8 post immunization (D) Endpoint titers against RBD for sera from Hartley guinea pigs immunized with RBD monomer and RBD g5.1 24mer after a single dose (E) Pseudo virus neutralization of sera from guinea pigs immunized with RBD monomer and RBD g5.1 24mer after a single dose (unpaired t-test vs. naïve: 5μg RBD g5. 1 24mer week 2 p=0.0140, week 3 p=0.0003, week 4 p=0.0146; 10μg RBD g5. 1 24mer: wk 2 p=0.0145, week 3 p=0.0016, week 4 p=0.0007. Unpaired t-test vs 10μg RBD to 5μg RBD g5. 1 24mer week 2 p=0.0142, week 3 p=0.0104; 10μg RBD g5. 1 24mer week 2 p=0.0002, week 3 p=0.0042, week 4 p=0.0304. (F) Endpoint binding titers against RBD for sera from Syrain Golden hamsters immunized with RBD monomer or RBD 48mer two times with two different doses (G) Neutralization of SARS-CoV-2 pseudovirus by sera from hamsters immunized with RBD monomer or RBD 48mer with two different doses. (unpaired two-tailed Student t-test: RBD 48mer 10 μg vs. RBD monomer 10 μg p = 0.0079, RBD 48mer 2 μg vs. RBD monomer 2 μg p = 0.0004) (H) Lung lavages from hamsters immunized with RBD monomer or RBD 48mer.

We assessed RBD g5.1 24mer in Hartley guinea pigs (n=6) to examine intradermal vaccine delivery at 0.5, 5 and 10μg in comparison to RBD monomer at 10μg. In contrast to the RBD monomer immunized group, we observed full seroconversion of RBD g5.1 24mer immunized animals at a dose of 5μg (Figure 6D). High levels of neutralizing antibodies were obtained in the 10μg dose group (ID_50_ of 1840) (Figure 6E). In a proof-of-concept study of RBD DLNPs prior to the development of RBD g5.1 24mer, we immunized Syrian Golden hamsters (n=5) twice 3 weeks apart with 2μg and 10 μg of RBD monomer and RBD 48mer (Figure 6F). The RBD nanoparticle immunized hamsters elicited higher antibody titers after both first and second doses and produced neutralizing antibodies unlike the responses in the RBD monomer vaccine groups (Figure 6G). To assess the biodistribution of anti-RBD IgG, we measured ultrafiltrated lung lavages and found antibodies only in the RBD nanoparticle groups (Figure 6H). In summary, we demonstrate that the DLNP vaccines provide enhanced immunogenicity in guinea pigs and hamsters.

## Discussion

New SARS-CoV-2 vaccines should (1) alleviate cold chain requirements for global vaccine distribution, (2) improve immunogenicity for certain populations, (3) increase efficacy with a single dose and (4) protect against emerging variants that reduce or evade current vaccine-induced immunity. We have demonstrated that advanced DNA formulation and delivery technology coupled with immune focused nanovaccines can provide a platform to address these translational obstacles for SARS-CoV-2 vaccines.

Viral glycan evolution results in antigenic changes with concomitant immune evasion. It has been observed that for influenza, humoral immunity becomes restricted over time due to glycan additions[61]. SARS-CoV-2 mutational variants may escape from antibody-mediated immunity. Glycan mutational variants may begin to circulate given their large impact on antibody recognition of virus. Here, we provide a map of possible glycan additions to the RBD of SARS-CoV-2 and their effect on a large series nAbs. Interestingly, we find a glycan at position 458, 369, 450 and 441 can bind to human ACE2, but strongly reduces binding to nAbs targeting the sites RBD-A, B, C and D respectively. We also show that addition of glycans to RBD-based nanovaccines can improve expression, assembly and immunogenicity.

The synthetic DNA platform employed in this study can be leveraged for generation of enhanced immunity, easier global distribution and rapid reformulation. New adaptive electroporation systems can improve uptake of DNA plasmid by 500x[62]. In stark contrast to complex recombinant protein and RNA-based product development, DNA-based production and purification are extremely easy due to availability of off the shelf commercial purification kits used widely in research laboratories. DNA vaccines are also much more chemically and thermally stable allowing storage at room temperature for long periods of time. These characteristics of the DNA platform allows for new vaccines to be developed at breakneck speed and distributed to resource limited settings around the globe.

A key finding in this study is the dose-sparing immunogenicity afforded by the nanoparticle designs. Most SARS-CoV-2 vaccines require at least two doses[63, 64]. DNA vaccines often require higher vaccine doses (25 μg in mice[41] and 5 mg in NHPs[65]) and/or advanced delivery devices to drive sufficient immunogenicity. Here, we observed strong immunogenicity and protection from bona fide SARS-CoV-2 challenge down to 1 μg. However, we did observe weight loss in 2/6 mice after challenge and low viral titers in 2/4 mice sacrificed four days after challenge. Increasing the dose to 10μg or by prime-boosting could improve on our results. In fact, our studies of DNA-launched nanoparticle vaccines in guinea pigs and hamsters demonstrated greater immunogenicity than RBD monomer at a low dose of 10μg.

In comparison to other RBD nanoparticle systems, we have demonstrated significant improvements. Recently, studies on two-component spike-based nanoparticle showed strong immunogenicity with sporadic pseudovirus neutralization 2 weeks post prime[66, 67]. After 2 doses of the i53-50 RBD nanoparticle vaccine, mice challenged with 1×10^5^ PFU of a mouse-adapted non-lethal virus were observed to have reduced viral replication[67]. SpyTag-coupled RBD nanoparticles induce binding but not neutralizing antibodies 2-weeks post prime[68–71]. An RBD-HR SpyTag nanoparticle was observed to induce immunity after 2 doses which after challenge with 4×10^4^ PFU authentic SARS-CoV-2 could reduce viral load in the lungs[72]. Here, the DNA-launched glycan modified RBDs could be genetically fused with *four* different nanoparticles scaffolds, the simple genetic fusion results in a single vaccine product that could induce binding and neutralizing antibodies 1 week post prime immunization and induce CD8+ Tcells. We created a more stringent test of immunity than most previous studies as we used authentic SARS-CoV-2 virus with 2.5-fold higher amount of virus (1×10^5^ PFU) in the challenge and a 10-fold more sensitive viral detection assay. Further, vaccines studied in this model mostly utilize a prime and boost to achieve protection (personal communication, Texas Biomed). In this model, our nanovaccines could induce immunity that reduced viral replication and completely protected from death at a low single dose of 5μg. From the data, there is an 82% chance of survival if mice have a live virus neutralization titer >100 prior to challenge. Given the protective threshold for neutralizing antibodies that we observe, we expect that the high levels of cross-reactivity neutralization to the B.1.1.7, B.1.351, P.1 SARS-CoV-2 variants generated by our nanovaccine would protect in a similar lethal challenge. In addition, the P.1 RBD g5.1 nanoparticle elicited high levels of cross-reactive antibodies which could be employed as a booster vaccine.

In conclusion, we have developed single-dose SARS-CoV-2 nanovaccine with a platform that can afford rapid pre-clinical reconfiguration to address variants of concern and for clinical translation.

## Methods

### Cloaking with Glycans Algorithm

The modeling started with RBD structure PDB id: 6M0J. GlycanTreeModeler(GTM) is a glycan modeling algorithm recently developed in Rosetta(unpublished). The Cloaking With Gycans (CWG) workflow utilizes GTM for selecting single glycan addition positions on target protein. All steps in CWG are summarized in a flowchart (Figure 1B). CWG begins with detecting native sequons and modeling all the native glycan structures using Man9GlcNAc2 glycans on the target protein. In the next stage, a model is made for the addition of a single glycan at each position. A given position in the protein is mutated to asparagine and the i+2 position is mutated into threonine or serine. The model with the lowest energy i+2 position is used for further evaluation. The Rosetta energy is computed for the resulting model. We filtered out positions if the total energy of the model corresponding to that position had a total energy > 5 Rosetta Energy Units (REU) more than the native structure. Next, the CWG algorithm builds Man9GlcNAc2 glycans on the mutated position and measures repulsive energy of engineered glycan between sugar-sugar and sugar-protein energy terms. We filtered out some positions based on structural criteria, such as avoiding the mutation of positions involved in disulfide bonds. Man9GlcNAc2 glycans were utilized for simplicity.

### Nanoparticle modeling

All nanoparticles were modeled with corresponding designed structures and linkers. Four nanoparticles were used in this study: IMX313P (PDB id: 4B0F), ferritin (PDB id: 3BVE), lumazine synthase (PDB id: 1HQK), and PcV (PDB id: 3J3I). Biological unit nanoparticle structure files were downloaded in CIF format. The termini of the monomeric RBDs were aligned to the termini of the nanoparticle, rotational and translational degrees of freedom were sampled to reduce clashing between RBDs and nanoparticles, extended linkers of various lengths were then aligned to fuse the nanoparticle and immunogen with simpleNanoparticleModeling from the MSL library as previously described[38].

### Protein expression and purification

Glycosylated RBDs: A gene encoding the amino acids 331-527 of the SARS-CoV-2 spike glycoprotein (PDB: 6M0J) was mutated at each position according to CWG. Nanoparticles were genetically fused to designed RBDs as described above. DNA encoding the variants were codon optimized for homo sapiens and cloned with a IgE secretion sequence into the pVAX vector. A 6xHisTag was added to the c-terminus of the RBD monomer variants. ExpiF293 cells were transfected with the pVAX plasmid vector either carrying the nanoparticles or the His-Tagged monomer transgene with PEI/Opti-MEM and harvested 6-7 days post transfection. The supernatants was first purified with affinity chromatography using the AKTA pure 25 purification system and IMAC Nickel column (HisTrap^™^ HP prepacked Column, Cytiva) for His-tagged monomers and gravity flow columns filled with Agarose bound Galnthus Nivalis Lectin beads (Vector Labs) for nanoparticles. The eluate fractions from the affinity chromatography were pooled, concentrated, and dialyzed into 1X PBS before being loaded onto the Size-Exclusion Chromatography (SEC) column for further purification with Superdex 200 Increase 10/300 GL column for the His-tagged monomers and the Superose 6 Increases 10/300 GL column for the nanoparticles. Fractions of interest were pooled and concentrated for characterization. For antibody production, heavy and light chains were encoded in pFUSEss-CHIg-hG1, and pFUSE2ss-CLIg-hk or pFUSEss-CLIg-hL2 respectively and were co-transfected in equal parts using ExpiFectamine^™^ 293 Transfection Kit(Gibco) according to manufacturer’s protocol. Antibodies were purified by affinity chromatography using the Protein A column (HiTrap^™^ MabSelect^™^ SuRe, Cytiva) and AKTA Pure 25 purification system.

### Western Blot

Samples were prepared with 13 μL supernatants of Expi293F cells transfected with RBD monomer plasmids or 0.65 μg of purified WT RBD in 1x PBS, NuPAGE LDS Sample Buffer (Novex), and NuPAGE Sample Reducing Agent (Novex) were denatured at 90°C for 10 minutes. Samples were loaded in a 4-12% SDS Bis-Tris gel for electrophoresis then transferred from the gel onto a PVDF membrane. The membrane was blocked with Intercept (PBS) Blocking Buffer (LI-COR) for >1 hour at ambient temperature then incubated with *** μg / protein gel of MonoRab anti-his tag C-term (Genscript) in Intercept T20 (PBS) Antibody Diluent (LI-COR) overnight at 4°C. The membrane was then incubated in a 1:10000 IRDye 800CW goat anti-rabbit IgG (LI-COR Biosciences) in Intercept T20 (PBS) Antibody Diluent (LI-COR) at room temperature for 1 h. Membranes were imaged with a LI-COR Odyssey CLx.

### ELISA

For in vitro characterization, high Binding, 96-well Flat-Bottom, Half-Area Microplate (Corning) were coated at 1 μg/mL 6x-His tag polyclonal antibody (Invitrogen) for >4 hours at ambient temperature and blocked ≥1 hour with 5% milk/1x PBS/0.01% Tween-20 at 4°C. RBD transfection supernatant or recombinant protein at 10 μg / mL was incubated for 1-2 hours at ambient temperature. Serial dilutions of antibodies were made according to affinity and incubated on plate for 1-2 hours at ambient temperature. Goat anti-Human IgG-Fc fragment cross-adsorbed antibody HRP conjugated (Bethyl Laboratories) secondary at a 1:10,000 dilution for 1 hour at ambient temperature. All dilutions except coating were performed in 5% milk/1x PBS/0.01% Tween-20 and plates were washed with 1x PBS/0.05% Tween-20 between steps. 1-Step^™^ Ultra TMB-ELISA Substrate Solution (Thermo Scientific) was incubated on the plate for 10 minutes in the dark and then quenched with 1 M H2SO4. Absorbance of samples at 570 nm was subtracted from 450 nm for each well and background of blank wells were subtracted from each well before analysis. Curves were analyzed in GraphPad Prism 8 with Sigmoidal, 4PL, X is concentration and AUC.

For serology, plates were coated with 1 μg/mL 6x-His tag polyclonal antibody (Invitrogen) in 1x PBS for 6 hours at ambient temperature and blocked overnight with 0.5% NCS/5% Goat Serum/5% Milk/0.2% PBS-T. 5x serial dilutions of sera were made starting at a 1:100 dilution and incubated on plate for 2 hours at 37 °C. For BL6, BALB/c, and K18 ACE2 mouse studies, goat anti-mouse IgG h+l HRP-tagged antibody (Bethyl Laboratories) diluted 1:20000. For the OmniMouse^®^ study, Peroxidase AffiniPure Goat Anti-Rat IgG (Jackson ImmunoResearch) at 1:10000, Peroxidase AffiniPure F(ab’)_2_ Fragment Goat Anti-Rat IgM, μ chain specific (Jackson ImmunoResearch) at 1:10000, Goat anti-Human Kappa Light Chain Antibody HRP Conjugated (Bethyl Laboratories) at 1:10000, Goat anti-Human Lambda Light Chain Antibody HRP Conjugated (Bethyl Laboratories) at 1:10000, and Goat anti-Mouse IgG-heavy and light chain Antibody HRP Conjugated (Bethyl Laboratories) at 1:20000, and Goat anti-guinea pig IgG whole molecule (Sigma) at 1:10,000 were used. Secondary antibodies were incubated on plates for 1 hr at RT. All dilutions except coating were performed in 1% NCS in 0.2% PBS-T and plates were washed with1x PBS/0.05% Tween-20 between steps. Plates were developed with 1 Step Ultra TMB substrate in the dark for 10 minutes for mouse studies and 15 minutes for guinea pig studies before being quenched with 1N H_2_SO_4_ and read using a BioTek Synergy 2 plate reader at an absorbance of 450 and 570nm.

Hamster serology was performed by directly coating 96-well flat bottom, half-area plates #3690 (Corning) with 25mL of 1 μg /mL of SARS-CoV-2 RBD (University of Texas, Austin) overnight at 4°C. Plates were blocked with 100uL of blocking buffer (3% BSA in 1 x PBS) for 1 hr at 37°C. Hamster sera was diluted to 1:16 dilution in diluent buffer (1% BSA in PBS) and an 11-point 1:3 serial dilution was done on the ELISA plate, with last column containing only dilution buffer as blank control. ELISA plates were incubated for 2 hr at 37°C with sera dilutions. Anti-Hamster HRP antibody (Sigma) was diluted in diluent buffer 1:10,000 and were incubated for 1 hr at room temperature. SureBlue TMB 1-Component Microwell Peroxidase Substrate (KPL) was added to the wells and plates were incubated for 6 minutes and then quenched with TMB Stop Solution (KPL). Absorbance was immediately read at 450 nm on Synergy HTX plate reader (BioTek). All volumes except blocking buffer was 25uL. Plates were washed 3 times with wash buffer (.05% Tween 20 in 1x PBS) between steps.

### Surface Plasmon Resonance

RBD-antibody kinetics experiments were performed with a Series S Sensor Protein A capture chip (Cytiva) on a Biacore 8k instrument (GE). The running buffer was HBS-EP (3 M sodium chloride/200 mM HEPES/60 mM EDTA/1.0% Tween 20 pH=7.6) (Teknova) with 0.1% (w/v) bovine serum albumin. Each experiment began with two start up cycles with 60 s of contact time and a flow rate of 50 μL/min. For analysis methods, approximately 200-300 RUs of IgG antibodies was captured on each flow cell at a flow rate of 10 μL/min for 60 seconds. WT RBD or glycan variants samples were 5x serial diluted from 1000 nM in running buffer and flowed across the chip after capture at a 50 μL/min rate. The experiment had a 120 second contact time phase and 600 seconds dissociation phase. Regeneration was performed with 10 mM glycine at pH=1.5 at a flow rate of 50 μL/min for 30 seconds after each cycle. Kinetic fits were analyzed with 1:1 fitting and run through a script to filter out results that had poor fitting, low max RUs compared to expected, and kon and koff constants that fell outside of the range of measurement. Experiments that were flagged as poor-quality fitting by this script were not further analyzed.

### Pseudovirus Neutralization Assay

HEK293T (CRL-3216) and CHO cells (CRL-12023: double check) were obtained from ATCC (Manassas, VA, USA). Cells were maintained in DMEM supplemented with 10% fetal bovine serum (FBS) and 1% penicillin-streptomycin (P/S) antibiotic at 37⍛C under 5% CO_2_ atmosphere. For luciferase-based virus pseudoneutralization assays, HEK293T cells were transfected to produce SARS-CoV-2 S containing pseudoviruses. Cells were seeded at 5 million cells onto T75 flasks and grown for 24 hours. Then, cells were treated with 48μL GeneJammer (Agilent 204130-21), 6μg S_IgE_deltaCterm19_plasmid (Genscript), and 6μg pNL4-3.luc.R-E- backbone (Aldevron) and incubated for 48 hours. For variant pseudoviruses, cells were similarly treated with GeneJammer and backbone with 6μg of S_SA_IgE_deltaCterm19, S_UK_IgE_deltaCterm19, or S_Brazil_IgE_deltaCterm19 plasmid. Transfection supernatants were then collected and supplemented with 12% FBS, sterile filtered, and stored at −80⍛C. Pseudovirus solutions were titered and dilution to working solutions set such that they yielded >215-fold greater relative luminescence units (RLS) than cells alone.

CHO cells expressing human ACE2 receptors (VCel-Wyb030) were obtained from Creative Biolabs (Shirley, NY). CHO-ACE2 cells were seeded at 10,000 cells/well in 96-well plates and incubated for 24 hours. Sera from vaccinated mice were heat inactivated at 56⍛C for 15 minutes. 3-fold serial dilutions starting at 1:20 dilutions in DMEM supplemented with 10% FBS and 1% P/S were performed on sample sera and incubated for 90 minutes at room temperature with SARS-CoV2 pseudovirus based on concentrations determined from titering described above. Media containing diluted sera and pseudovirus were then applied to CHO-ACE2 cells. After 72 hours of incubation, cells were developed using BriteLite plus luminescence reporter system (Perkin Elmer 6066769) and signal measured using a plate reader (Biotek Synergy). Percent neutralization was calculated based on virus only positive control signal with background subtraction of cells only negative controls. ID_50_ values were calculated using GraphPad Prism v8.0 nonlinear curve fitting with constraint Hill Slope < 0.

### SARS-CoV-2 culture, titer, and neutralization assay

SARS-Related Coronavirus 2, Isolate USA-WA1/2020, NR-52281 was deposited by the Centers for Disease Control and Prevention and obtained through BEI Resources, NIAID, NIH. All work with it was performed in the BSL-3 facility at the Wistar Institute. Vero cells (ATCC CCL-81) were maintained in antibiotic-free Dulbecco’s modified Eagle’s medium (DMEM) supplemented with 10% fetal bovine serum (FBS). To grow a stock of virus, 3 million Vero cells were seeded in a T-75 flask for overnight incubation (37⍰C, 5% CO2). The cells were inoculated the next day with 0.01 MOI virus in DMEM. Culture supernatant was harvested 3 days post infection, aliquoted, and stored at −80°C. For titering the virus stock, Vero cells were seeded in DMEM with 1% FBS at 20,000 cells/well in 96 well flat bottom plates for overnight incubation (37⍰C, 5% CO2). The USA-WA1/2020 virus stock was serially diluted in DMEM with 1% FBS and transferred in replicates of 8 to the previously seeded Vero cells. Five days post infection individual wells were scored positive or negative for the presence of cytopathic effect (CPE) by examination under a light microscope. The virus titer (TCID50/ml) was calculated using the Reed-Munch method and the Microsoft Excel based calculator published by Lei et al[73] For neutralization assays, Vero cells were seeded in DMEM with 1% FBS at 20,000 cells/well in 96 well flat bottom plates for overnight incubation (37C, 5% CO_2_). Serum samples were heat inactivated at 56°C for 30 minutes. Serum samples were then serially diluted in DMEM with 1% FBS and 1% penicillin/streptomycin and incubated for one hour at room temperature with 300 TCID50/ml USA-WA1/2020. The serum-virus mixture was then transferred in triplicate to the previously seeded Vero cells. Five days post infection, individual wells were scored positive or negative for the presence of CPE and neutralization titers were calculated using the Reed-Munch method and a modified version of the Microsoft Excel based calculator published by Lei et al[73].

### Animal Studies

C57BL/6, BALBc, and K18-hACE2 mice were obtained from Charles River Laboratories (Malvern, PA) and The Jackson Laboratory (Bar Harbor, ME). Omni Mouse^®^ for human antibody studies were obtained from Ligand Pharmaceuticals Incorporated (San Diego, CA). All studies were performed in accordance with Wistar Institutional Animal Care and Use Committees under approved animal protocols. All animals were housed in the Wistar animal facility in ventilated cages and given free access to food and water. For the lethal challenge study, Texas Biomed were blinded to identity of vaccination groups and weight loss cutoff for euthanasia was 20%. Intramuscular injection with electroporation and sample collection. Plasmids were administered intramuscularly in 30uL water into the tibialis anterior muscle. Electroporation was then performed using CELLECTRA EP delivery platform consisting of two pulse sets at 0.2 Amps at a 3 second interval. Each pulse set consists of two 52 ms pulses with 198 ms delay. At specified time points, blood was collected via submandibular vein puncture and centrifuged for 10 min at 15000 rpm to obtain sera. For cellular responses, mice were euthanized under CO_2_ overdose. Spleens were collected into cold RPMI media supplemented with 10% FBS and 1% P/S.

Female Hartley guinea pigs (8 weeks old, Elm Hill Labs, Chelmsford MA) were housed at Acculab (San Diego CA). On day 0 and day 28 animals were anaesthetized with isoflurane vapor and received intradermal Mantoux injections of 100 μL 10, 5 or 0.5 μg pDNA immediately followed by CELLECTRA-3P electroporation. The CELLECTRA^®^ EP delivery consists of two sets of pulses with 0.2 Amp constant current. Second pulse set is delayed 3 seconds. Within each set there are two 52 ms pulses with a 198 ms delay between the pulses. Serum samples were collected by jugular or saphenous blood collection throughout the study on days 0, 7, 14, 21, 28 and 42. Whole blood samples to process PBMCs for cellular assay were collected from the jugular vein on days 14 and 42. All animals were housed in the animal facility at Acculab Life Sciences (San Diego, CA). All animal protocols were approved by Acculab Institutional Animal Care and Use Committees (IACUC).

Golden Syrian hamsters (8 weeks old, Envigo, Indianapolis, IN) were housed at Acculab (San Diego, CA). Hamsters received intramuscular (IM) injections of 60 μL of 2 or 10μg pDNA formulation into the tibialis anterior muscle immediately followed by electroporation with the CELLECTRA-3P device under Isoflurane vapor anesthesia at day 0 and day 21. The CELLECTRA^®^ EP delivery consists of two sets of pulses with 0.2 Amp constant current. Second pulse set is delayed 4 s. Within each set there are two 52 ms pulses with a 198 ms delay between the pulses. Serum samples were collected at indicated timepoints via saphenous vein blood collection throughout the experiment. All animals were housed in the animal facility at Acculab Life Sciences (San Diego, CA). All animal protocols were approved by Acculab Institutional Animal Care and Use Committees.

In-vivo study was concluded with terminal blood, lung lavage and nasal wash collection. Lavage buffer was prepared as PBS containing 100uM EDTA, 0.05% Sodium Azide, 0.05% Tween-20 and Protease Inhibitor. Hamsters were euthanized by jugular exsanguination with intraperitoneal (IP) injection of 86.7mg/kg pentobarbital sodium or overdose Isoflurane gas inhalation. Euthanized hamster was placed in supine position and skin was disinfected using 70% Isopropyl alcohol. A longitudinal cut using scissors and blunt dissection along the midline of the neck was performed to expose the trachea. An opening into the exposed trachea was created by making a transverse, semilunar cut using #11 blade.

To collect nasal wash an 18ga blunt end needle was inserted toward the nose and gently proceeded upwards until reaching the nasal palate. A syringe filled with 1.5mL lavage buffer was connected to the blunt end needle and correct placement was tested by dispensing a small amount through the hamster’s nares. The entire volume of lavage fluid was rapidly dispensed and collected directly from the nares into a 5.0mL Eppendorf tube.

To collect bronchioalveolar lavage (BAL), an 18ga blunt end needle, attached to a three-way stopcock and primed with lavage buffer (approximately 0.5mL) to eliminate empty airspace, was inserted forward until just prior to the tracheal bifurcation into the lungs. The blunt end needle was secured in the trachea with a silk 2-0 tie. A 3mL receiver syringe and a 10mL syringe filled with 9mL of lavage buffer was connected to the blunt end needle via the three-way stopcock. The lungs were rinsed three times (3mL each time) with a total of 9mL lavage buffer. Typically, 50% of lavage buffer was recovered.

### Hamster biodistribution

Lung lavage and nasal wash samples were ultrafiltrated using a 2mL 100kDa cut-off ultrafiltration device (Millipore, Burlington MA) spinning 1mL BAL or NW for 15min at 4000g. Ultrafiltrated BAL was diluted 1:6 and nasal wash was diluted 1:4 in ELISA dilution buffer and following washes and blocking as described in the ELISA section added to half area assay plates (Costar) coated with 25μL/well of 1 μg/mL SARS-CoV-2 RBD (Sinobiological) in dilution buffer overnight at 4C. BAL and NW samples were tested at a 7-step 1:2 serial dilution.

### Negative-stain electron microscopy

Purified RBD g5.1 nanoparticle was dialyzed into 20 mM HEPES buffer, 0.15M NaCl, pH 7.4. A total of 3 μL of purified proteins was adsorbed onto glow discharged carbon-coated Cu400 EM grids. The grids were then stained with 3 μL of 2% uranyl acetate, blotted, and stained again with 3 μL of the stain followed by a final blot. Image collection and data processing was performed on a FEI Tecnai T12 microscope equipped with Oneview Gatan camera at 90 450× magnification at the camera and a pixel size of 1.66 Å.

### Cryo electron microscopy

Cryo-EM vitrification was obtained in a Vitrobot Mark IV robot (FEI). Four μL of purified RBD g5.1 24mer nanoparticles in 1xPBS were deposited on a glow-discharged holey carbon grid (C-flat 1.2/1.3, 300 mesh; Protochips). Excess liquid was blotted away followed by immediate plunging into liquid ethane cooled by liquid nitrogen. The vitrified specimen was then introduced into an FEI Talos Arctica electron microscope (FEI). Automated data collection was performed in EPU (FEI) and 640 movie micrographs were recorded with a Falcon 3 camera (FEI) at 150,000x magnification corresponding to an image pixel size of 0.97Å on the object scale. Each movie micrograph comprised 50 frames, each frame was exposed with a dose of ~1 e^-^/Å^2^. Data processing was performed in Relion v3.1.2[74]. Movie micrograph frame alignment, spectral signal weighing and summation was followed by CTF modeling (CTFFIND4[75]). Candidate molecular projection images were identified with Relion LoG picking (~271,000). Image windows corresponding to the candidate molecular projection image coordinates were extracted and binned by a factor of 2. The extracted binned data was subjected to 2D classification. Manual inspection of class averages led to identification of 93,348 molecular projection images selected for further data processing. Molecular projections were re-extracted unbinned from the summed micrographs and iterative Euler angular reconstitution and 3D object reconstruction was performed with a low-resolution ferritin density map as initial seed. 3D refinement was performed both asymmetrically (FSC 0.143 resolution 3.98Å) and under the assumption of octahedral symmetry (FSC 0.143 resolution 3.42Å). Since our objective was to map the attachment sites of the RBDs to the ferritin cage, we made no efforts to improve ferritin particle alignment in our refinement strategy for the present manuscript.

### ELISpot assay

Spleens from immunized mice were processed by a tissue stomacher, and red blood cells were then lysed by ACK buffer (Thermo Fisher Scientific). Single cell suspension was counted, and 2 x 10^5^ splenocytes were plated into each well of the Mouse IFN-γ ELISpotPLUS plates (MabTech). The splenocytes were stimulated for 20 hours at 37°C with RBD peptides (15-mer peptides overlapping by 9 amino acid spanning the RBD of SARS-CoV-2 spike protein, GenScript), at 5μg/mL of each peptide in RPMI + 10% FBS (R10). The spots were developed according to manufacturer’s instructions. R10 and cell stimulation cocktails (Invitrogen) were used for negative and positive controls, respectively. Spots were scanned and quantified by ImmunoSpot CTL reader. Spot-forming unit (SFU) per million cells was calculated by subtracting the negative control wells.

### Intracellular cytokine staining and Flow cytometry

Splenocytes were processed as described in the previous section and stimulated with RBD peptides for 5 hours at 37°C with protein transport inhibitor (Invitrogen) and anti-mouse CD107a-FITC antibody (BioLegend). Cell stimulation cocktail and R10, with protein transport inhibitor, were used as positive and negative controls, respectively. After stimulation, cells were stained with Live/Dead violet (Invitrogen) for viability. Anti-mouse CD4-BV510, CD8-APC-Cy7, CD44-A700, and CD62L-BV711 antibodies were used for surface staining and CD3e-PE-Cy5, IFN-γ-APC, and TNF-α-BV605 (all from BioLegend) were used for intracellular staining. The samples were run on an 18-color LSRII flow cytometer (BD Biosciences) and analyzed by FlowJo software.

### Competition assay

96-well Flat-Bottom Half-Area plates (Corning) were coated at room temperature for 8 hours with 1 μg/mL 6x-His tag polyclonal antibody (PA1-983B, ThermoFisher), followed by overnight blocking with blocking buffer containing 5% milk/1x PBS/0.01% Tween-20 at 4°C. The plates were then incubated with RBD at 1 μg/mL at room temperature for 1-2 hours. Mouse Sera (BALB/c, terminal bleeds, week 6, n=5) either immunized with RBD-WT or RBD-gPenta was serially diluted 3-fold starting at 1:20 with dilution buffer (5% milk/1x PBS/0.01% Tween-20) was added to the plate and incubated at room temperature for 1-2 hours. Plates were then washed and incubated at room temperature for 1 hour with ACE2-IgHu at a constant concentration of 0.06μg/mL diluted with the dilution buffer. After being washed, the plates were further incubated at room temperature for 1 hour with goat-anti human IgG-Fc fragment cross-adsorbed Ab (A80-340P; Bethyl Laboratories) at a 1: 10,000 dilution, followed by addition of TMB substrates (ThermoFisher), and then quenched with 1M H_2_SO_4_. Absorbances at 450nm and 570nm were recorded with a BioTek plate reader. Four washes were performed between every incubation step using PBS and 0.05% Tween-20. The assay was performed in triplicates. The average absorbance of the lowest dilutions with saturating ACE2 signals was calculated to get a maximum ACE2 binding and no blocking. Each average absorbance value was subtracted from the maximum to get an ACE2 blocking curve. The blocking titer is defined as the reciprocal of the highest dilution where two consecutive dilutions have readings below zero. The maximum area under the curve is determined by calculating the Area Under the Curve (AUC) of full ACE2 binding without the competitor. The AUC of the competitor is then subtracted from the maximum AUC which provides the area between the two curves (blocking area) and is a measure of ACE2 blocking. The fraction ACE2 blocking is defined as the fraction of the blocking area to the maximum AUC.

## Figure Legends

**Supplementary Figure 1:**
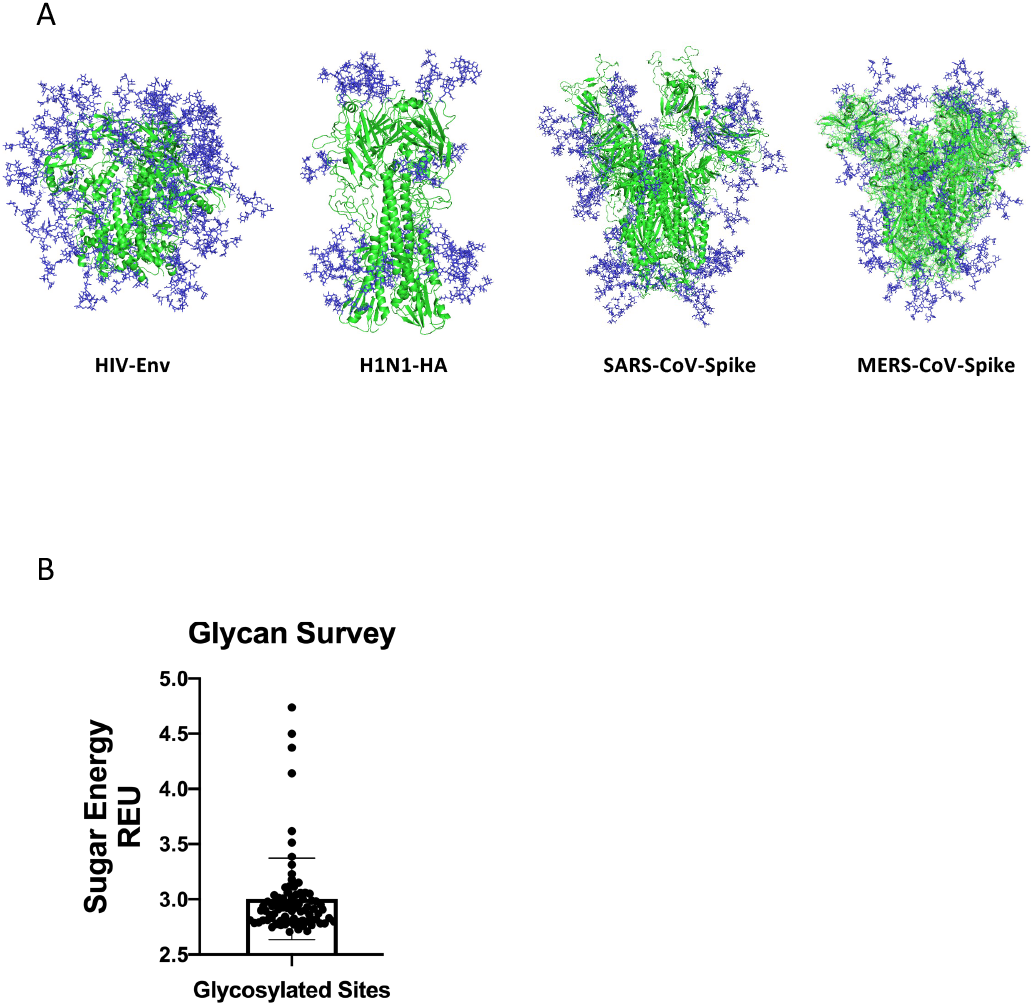
Modeling and Survey of native glycans on human virus proteins. (A) Native glycans were modeled on PNGS using our modified Rosetta GlycanTreeModeler script on four human viral glycoproteins: envelope of HIV, hemagglutinin of H1N1, and Spike proteins of SARS-CoV and MERS-CoV. (B) Repulsive glycan energy of individual modeled glycans on each native PNGS sites were surveyed and a cut-off value of 5.0 (REU) is used to include all possible native glycosylation scenarios.

**Supplementary Figure 2:**
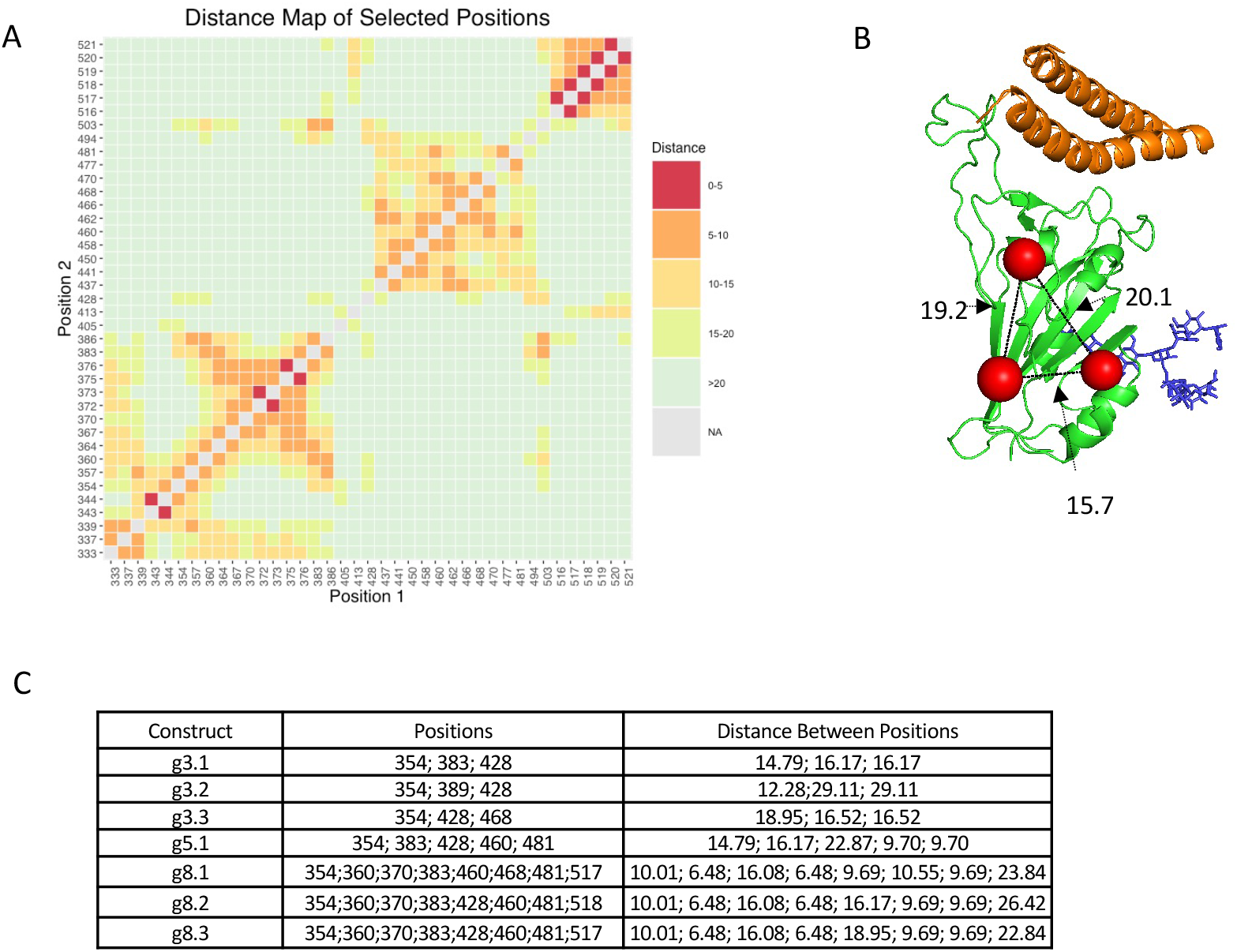
Rationale for generating glycan combinations. (A) Distance map of all residue pairs on native RBD to approximate distances between two engineered glycans. Distance between the geometric center of each residue of any residue pair on RBD is calculated using PyRosetta script. The distance information is subsequently visualized as a Distance Heatmap in R. Only glycans that are 10-20 Angstroms away from each other can be selected for a combination. (B) An example of distance measurements for a combination of three glycan additions where the residue of added glycans (red spheres) is on RBD (green) bound to two helices of ACE2 (orange) with one native glycan (blue). Distances between each engineered glycan is shown in black dash and labeled with the distance value in Angstroms. (C) A table summary of all combinations made in this study with glycan addition positions and distance between engineered glycans.

**Supplementary Figure 3:**
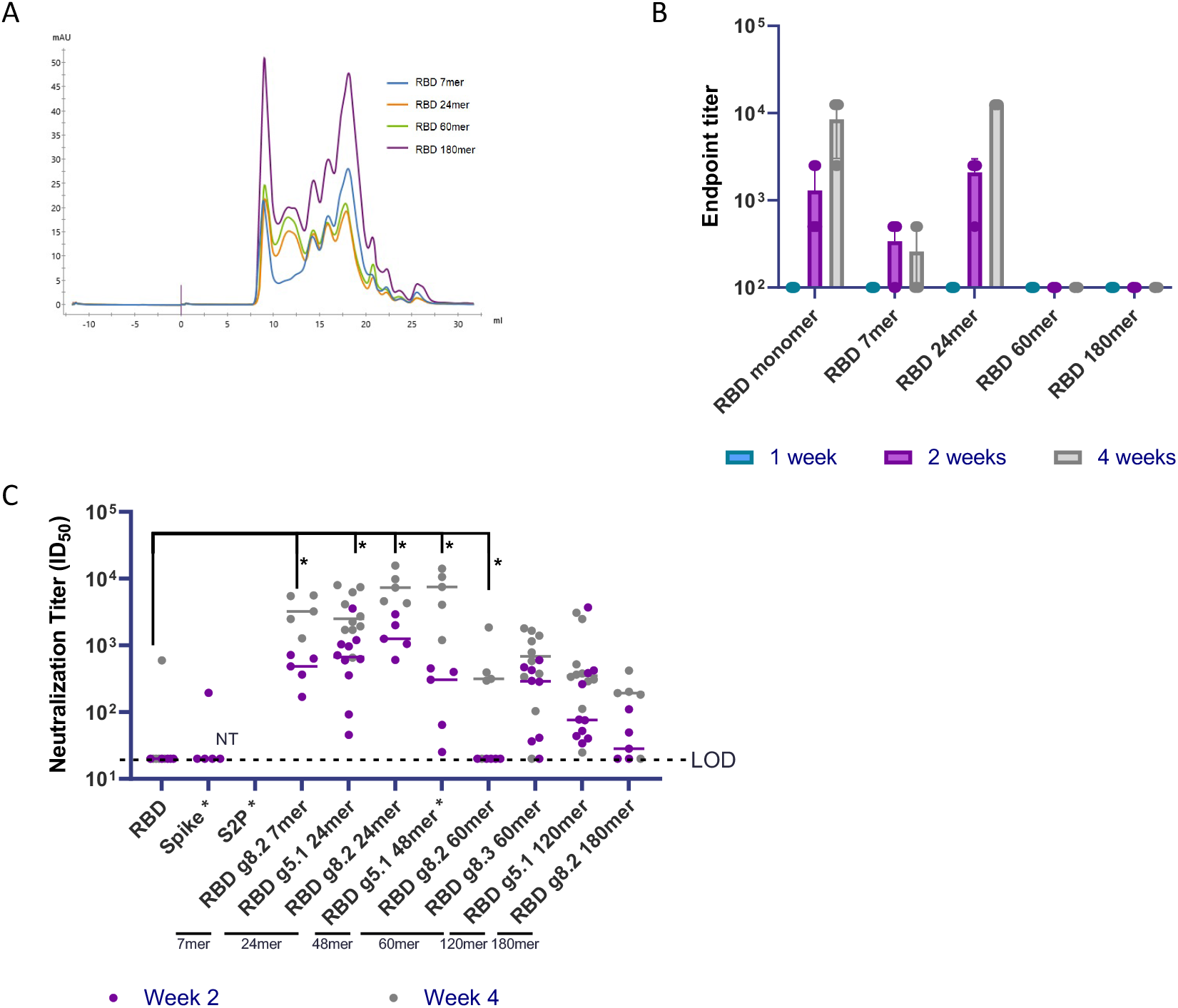
WT RBD nanoparticles and pseudovirus neutralization of expanded groups (n=5) in BALB/c mice. (A) Size exclusion chromatograms for WT RBD nanoparticles. (B) Endpoint titers from binding ELISA to RBD of immunizations with 2μg of DNA-launched WT RBD nanoparticles. (C) Pseudovirus neutralization of BALB/c mice immunized (n=5 or 10) with 2 μg immune focused DNA-launched nanoparticles. * p < 0.05 (Two-way ANOVAs vs. RBD: RBD g8.2 7mer p = 0.0111, RBD g5.1 24mer p = 0.0205, RBD g8.2 24mer p = 0.0199, RBD g8.2 60mer p = 0.0135).

**Supplementary Figure 4:**
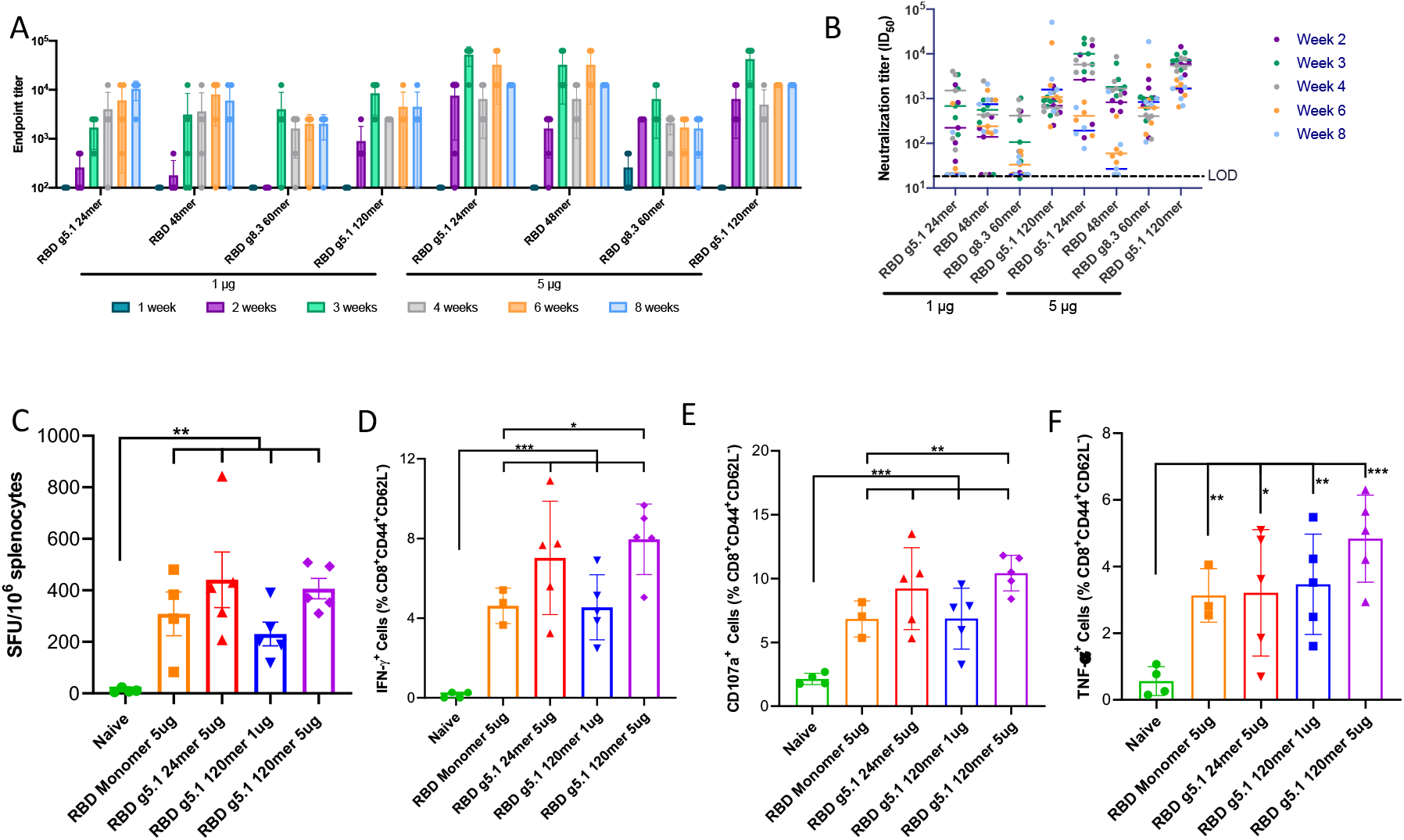
Immunogenicity of RBD nanovaccines in C57BL/6 mice. Endpoint titers (A) and pseudo virus neutralization ID50s (B) for C57BL/6 mice immunized with 1 μg or 5 μg of four selected RBD nanovaccines. (C) IFN-γ ELISpot assay with splenocytes from mice immunized with RBD monomer, RBD g5.1 24mer, and RBD g5.1 120mer vaccines, or the naive. Intracellular staining of IFN-γ (D), surface staining of CD107a (E), and intracellular staining of TNFα (F) of effector memory CD8+ CD44+ CD62-T cells from splenocytes. Error bars indicate means ± SD (n = 3 - 5 mice/group). Splenocytes were stimulated by native RBD peptides in B, C, and D. ((C) unpaired two-tailed Student t-tests vs. naïve: RBD monomer p = 0.0063, RBD g5.1 24mer p = 0.0049, RBD g5.1 120mer 1 μg p < 0.0019, RBD g5.1 120mer 5 μg p < 0.0001). ((D) unpaired two-tailed Student t-tests vs. naïve: RBD monomer p < 0.0001, RBD g5.1 24mer p = 0.0010, RBD g5.1 120mer 1 μg p = 0.0005, RBD g5.1 120mer 5 μg p < 0.0001; vs. RBD Monomer: RBD g5.1 120mer 5 μg p = 0.0123). ((E) unpaired two-tailed Student t-tests vs. naïve: RBD monomer p = 0.0007, RBD g5.1 24mer p = 0.0018, RBD g5.1 120mer 1 μg p = 0.0031, RBD g5.1 120mer 5 μg p < 0.0001; vs. RBD Monomer: RBD g5.1 120mer 5 μg p = 0.0063). ((F) unpaired two-tailed Student t-tests vs. naïve: RBD monomer p = 0.0013, RBD g5.1 24mer p = 0.0153, RBD g5.1 120mer 1 μg p = 0.0038, RBD g5.1 120mer 5 μg p = 0.0002).

**Supplementary Figure 5:**
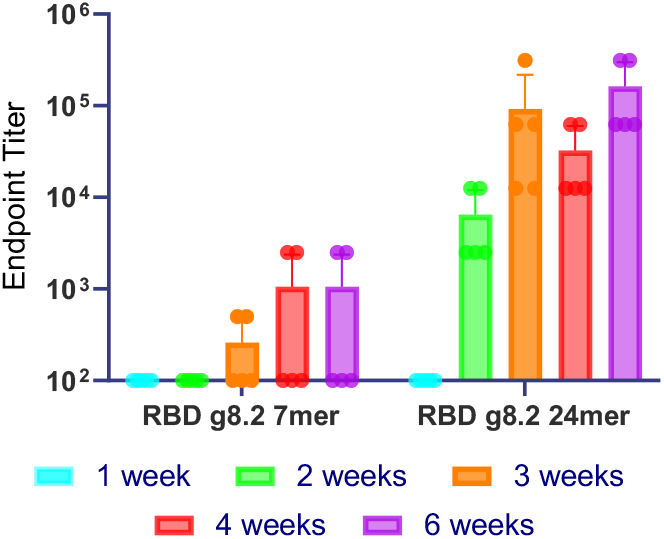
Protein nanoparticle immunization. Endpoint titers of BALB/c mice immunized SC with 10 μg of RBD g8.2 7mer and 24mer protein co-formulated with RIBI adjuvant.

**Supplementary Figure 6:**
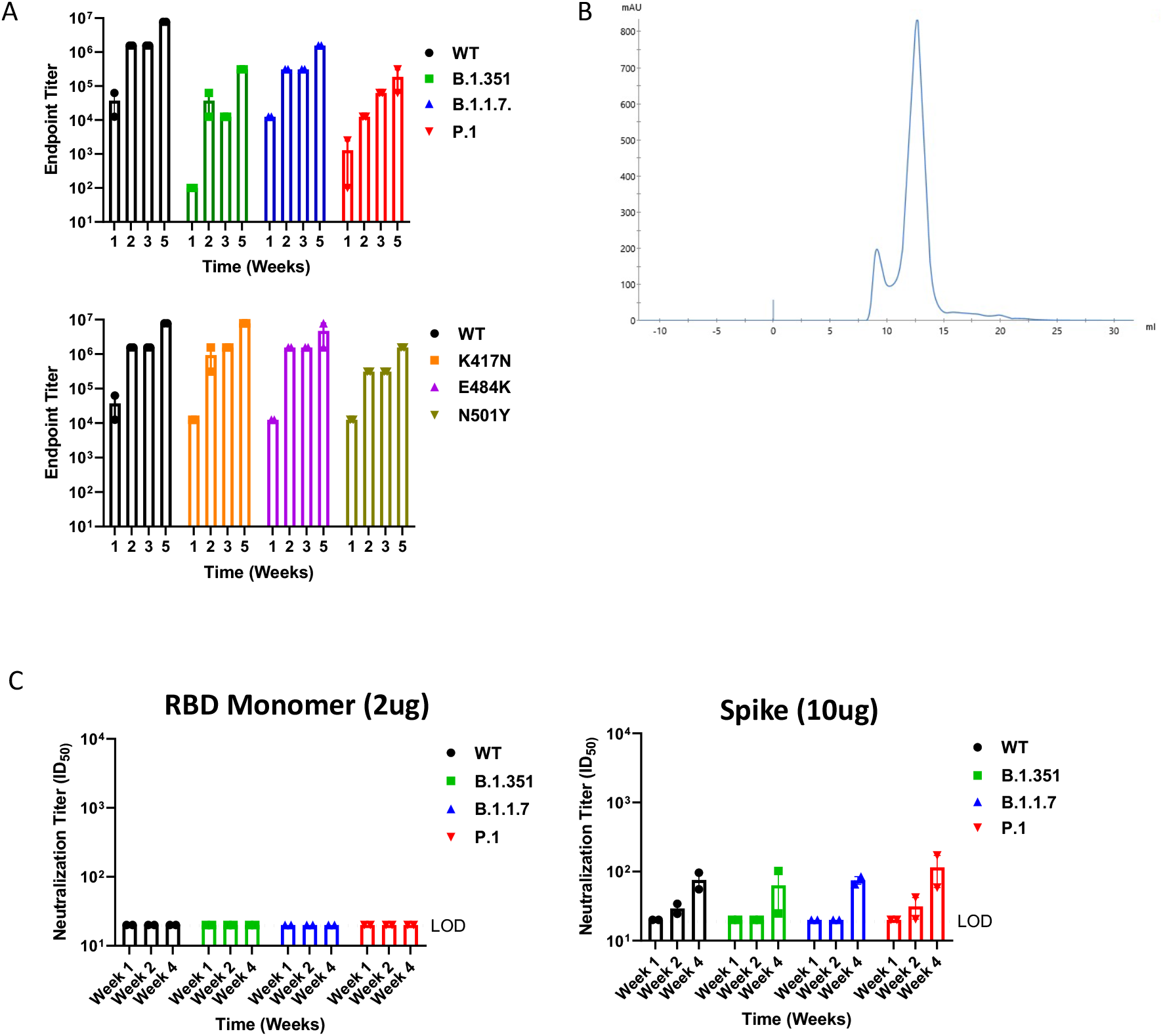
Emerging variants of concern. (A) Endpoint titers of BALB/c mice immunized with 5μg of RBD g5.1 24mer (WT RBD) binding to WT, B.1.351, B.1.1.7., and P.1 RBDs and individual mutation RBDs. (B) SEC trace of P.1 RBD g5.1 24mer. (C) Comparison of neutralization of BALB/c mice immunized with RBD monomer 2 μg and Spike 10 μg against variant pseudoviruses.

**Supplementary Figure 7:**
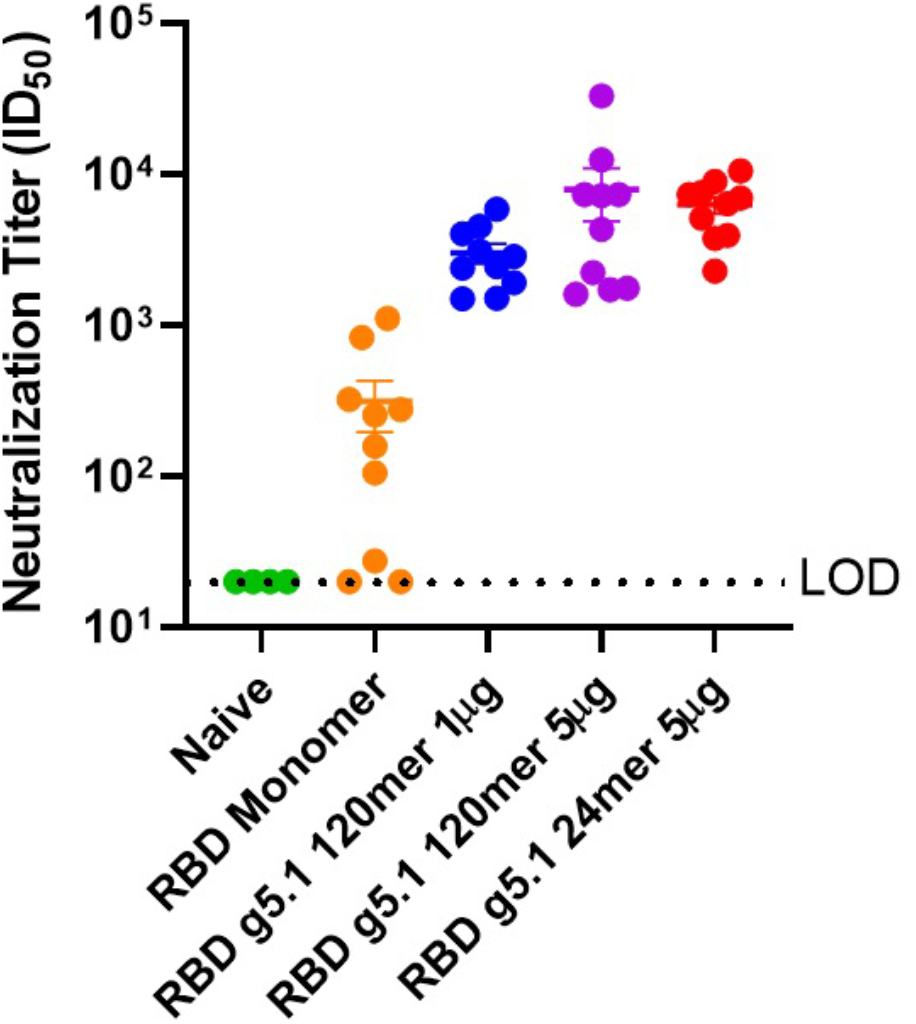
Pre-challenge pseudo virus neutralization of K18 hACE2 mice. Pseudo virus neutralization of week 3 sera from K18 hACE2 mice immunized for challenge study.

**Supplementary Figure 8:**
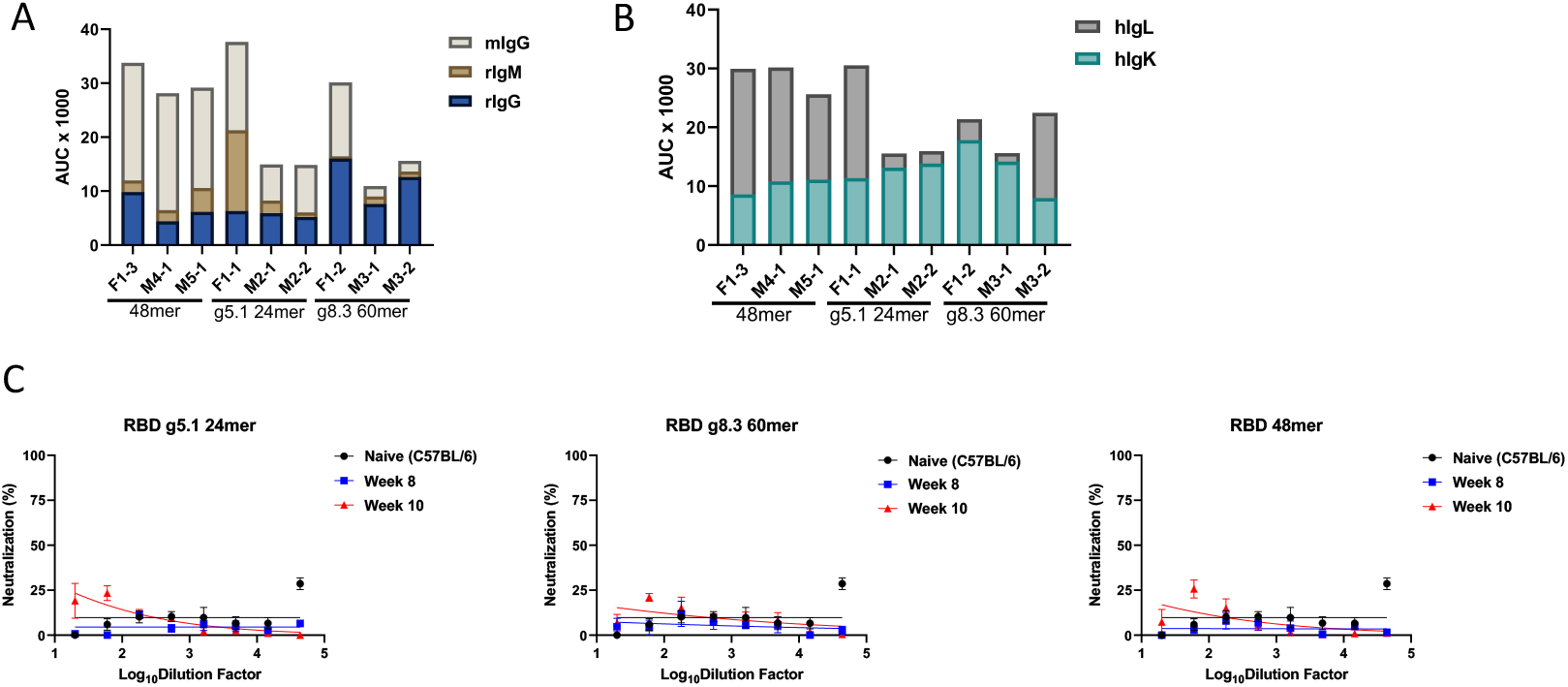
Additional Omni mice serology. (A) AUC of binding ELISAs from Omni mice immunized with DNA-launched RBD nanoparticles with mIgG, rIgM, and rIgG breakdown. (B) AUC of binding ELISAs from Omni mice with hIgL and hIgK breakdown. (C) Murine leukemia virus (MLV) neutralization of Omni mice immunized with DNA-launched RBD nanoparticles demonstrate no nonspecific neutralization.

**Supplementary Figure 9:**
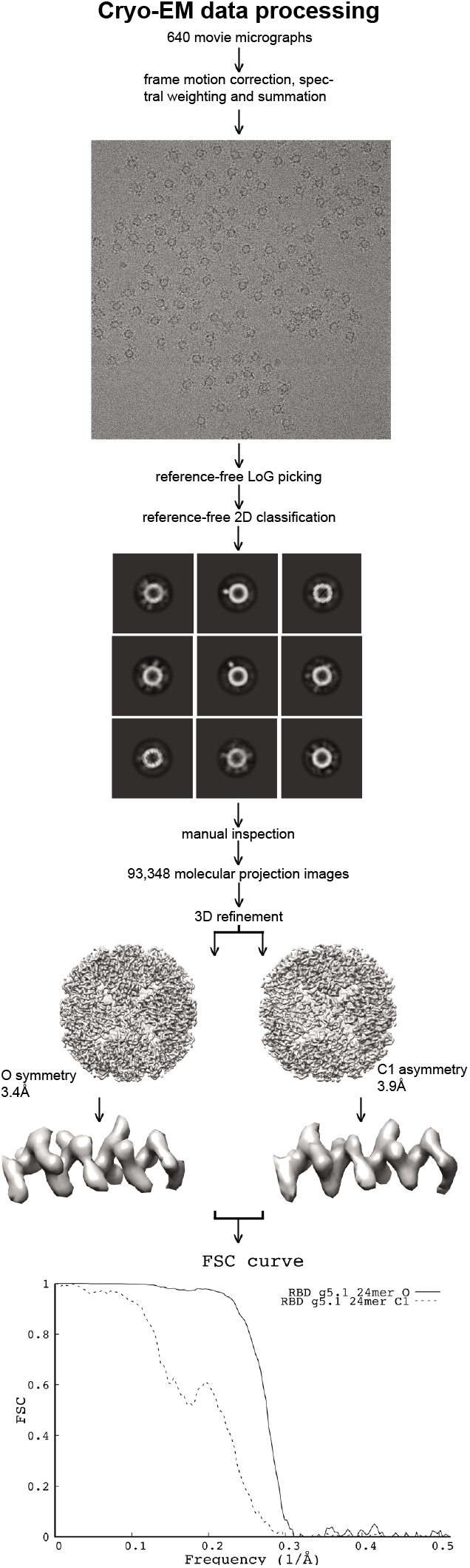
Cryo-EM of RBD g5.1 24mer immunogen. Cryo-EM data processing followed standard routines. 3D reconstruction was performed under assumption of octahedral symmetry as well as asymmetrically. The two resulting density maps demonstrate flexible linker attachment points at low density threshold at identical places. Final resolutions were 3.4Å and 3.9Å, respectively as can be confirmed by visual inspection of the density close-ups. Density for RBDs is disordered due to the inherent flexible linker in our immunogen design.

**Table 1.**
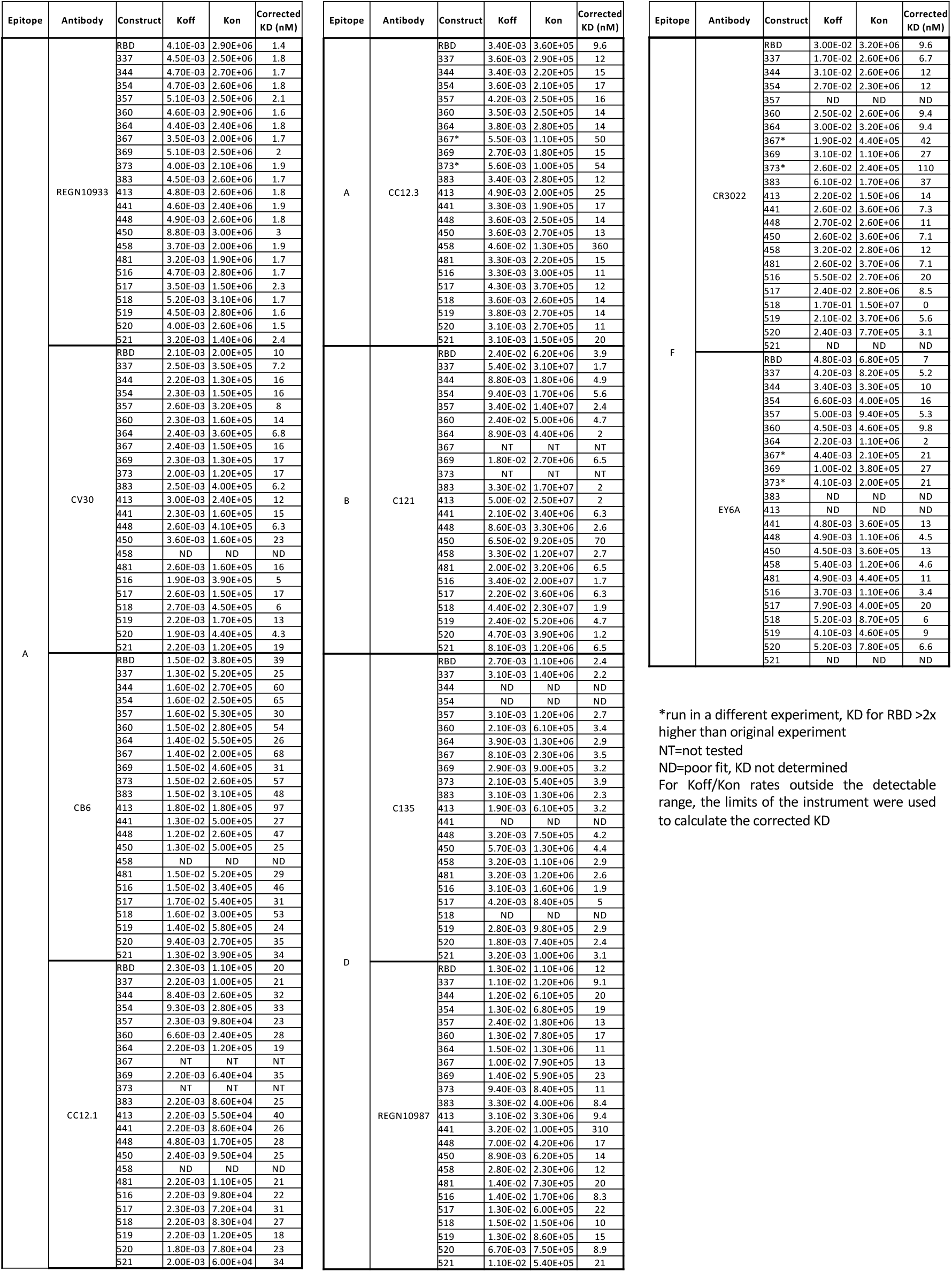
Kinetic constants for RBD-antibody interactions modeled well by 1:1. Langmuir fitting.

## Acknowledgments

The authors would like to thank the The Wistar Institute Core facilities for providing care to the animals. We would like to acknowledge Dr. Jason S. McLellan for providing reagents for hamster serology. We would also like to thank Dr. Jared Adolf-Bryfogle for generously contributing GTM code to Rosetta for use in this project.

## Author Contributions

K.M.K., Y.W. and D.W.K. designed immunogens. K.M.K, K.L., Z.X., S.N.W., X.Z., N.C., N.T., M.P., J.P., E.L.R., D.F., C.I. D.B.W and D.W.K. planned experiments. K.M.K., K.L., Z.X., S.N.W., X.Z., N.C., N.T., M.P., J.P., J.D., A.M., E.L.R. and D.F conducted experiments. K.S., K.E.B., L.H., and T.S. contributed resources for lethal challenge study. C.I. contributed resources for the human antibody transgenic mouse study. K.M.K., K.L., Z.X., S.W., X.Z., N.C., N.T., M.P., J.P., E.L.R., J.D., A.M., D.F. and D.W.K. analyzed the data. K.M.K. and D.W.K. wrote the article. K.M.K, K.L., Y.W., S.N.W., N.C., J.D., A.M., J.P., D.B.W., and D.W.K. edited the article.

